# A fast neutron mutant resource for *Ipomoea purpurea* reveals dose-dependent genomic and phenotypic variation

**DOI:** 10.64898/2026.10.19.760188

**Authors:** M. Inam Jameel, Madisyn Guza, Meredith Cobb, Ethan A. Baldwin, James H. Leebens-Mack, Wayne A. Parrott, Regina S. Baucom

## Abstract

A central goal in evolutionary biology is to understand the genomic basis of phenotypic traits, yet this pursuit is often constrained by a lack of accessible phenotypic and genetic variation, particularly in non-model organisms. Fast neutron irradiation offers a powerful approach for generating such variation, as it does not require prior genomic resources, established transformation systems, or reference mutant collections. Here, we describe the development and characterization of the first fast neutron mutant library in *Ipomoea purpurea* (common morning glory), a noxious agricultural weed and established model system in evolutionary ecology. We exposed an *I. purpurea* reference genotype to three levels of irradiation and developed 1,043 lines third generation lines and characterized the phenotypic and genetic variation in a subset of 200. Phenotypic screens identified seven mutant lines exhibiting eleven distinct phenotypes, and revealed higher phenotypic variance for anther-stigma distance, flower number, and pollen count relative to a non-irradiated control line. We further identified genetic variation across eight phenotypic traits among the mutant lines. Together, these results show that fast neutron mutagenesis generates novel phenotypic and genetic variation in ecologically relevant traits, offering the research community a new genomic resource for an established model in evolutionary ecology.

## Introduction

A central goal in evolutionary biology is to understand the genomic basis of phenotypic traits, yet progress toward this goal is often limited by a lack of accessible phenotypic and genetic variation, particularly in non-model organisms. Induced mutagenesis is a well-established technique used by plant researchers to generate novel variation and study gene function (Koornneef et al. 1982; Chen et al. 2022), offering one route to address this shortcoming. Among mutagenesis approaches, fast neutron mutagenesis is particularly versatile, inducing a broad spectrum of mutations–from single nucleotide polymorphisms to insertions and deletions–primarily through the generation of double strand breaks (Li et al. 2016; Wyant et al. 2022; Yuan et al. 2024). This flexibility has made fast neutron populations valuable across a range of biological contexts, including validating genes underlying nodulation trait variation in legumes (Men et al. 2002; Hoffmann et al. 2007; Bolon et al. 2011) and examining variation in salt tolerance in wheat (Hanafy and Mohamed 2014).

Despite this versatility, fast neutron populations have been developed predominantly in crop and laboratory model systems (Bolon et al. 2011; Kumawat et al. 2019; Wyant et al. 2022; Yuan et al. 2024), reflecting historical emphasis on pragmatic questions in crop improvement. However, the utility of these resources extends beyond applied research programs. Other disciplines such as ecology and evolution stand to benefit equally from these resources, particularly for identifying the genetic basis of adaptive traits (Bomblies and Peichel 2022). Fast neutron resources remain largely absent from weed biology and evolutionary ecology specifically, signaling a disparity between the technique’s demonstrated versatility and its application to questions of natural selection and ecologically and evolutionarily relevant trait distributions. Developing such resources in an evolutionary ecology model species would help close this gap.

Many ecologically and agriculturally important traits are quantitative, governed by multiple genes of small, additive effect rather than single loci of large effect. Because mutational effects in such traits are distributed across the genome, induced mutagenesis offers a means of directly influencing trait variance (Stearns and Fenster 2016), an approach with roots in classic radiation biology (Mertens and Burdick 1957; Oka et al. 1958; Rawlings et al. 1958; Daly 1973), and more recently demonstrated in fast neutron and T-DNA insertion populations, where induced mutations generate extreme phenotypes for quantitative traits (Bolon et al. 2011; Rutter et al. 2019). Examining variance across a range of irradiation doses further allows for the detection of dose-dependent effects on quantitative variation. This variance-based approach complements existing genomic methods, which are well-suited to identifying discrete causal mutations but less suited to characterizing their impact on continuous trait distributions.

*Ipomoea purpurea* (Convolvulaceae), or the common morning glory, is a weedy annual vine that has served as a productive study system for questions in ecology and evolution, including herbicide resistance (Baucom et al. 2011; Kuester et al. 2015; Van Etten et al. 2016; Gupta et al. 2023), mating system evolution (Chang and Rausher 1998; Kuester et al. 2017), plant-pollinator (García et al. 2023) and plant-herbivore interactions (Zhang and Baucom 2025). Beyond its established role in evolutionary ecology, *I. purpurea* offers several practical advantages for developing a mutant resource: high seed production, reliable germination, and a short, six-week generation time under short-day conditions. These attributes are further supported by an annotated draft genome assembly (Gupta et al. 2023), which provides the base scaffolding for interpreting the genomic and functional consequences of induced mutations. Combined, these ecological, practical, and genomic resources position *I. purpurea* as a system uniquely suited to linking induced mutations to ecologically and evolutionarily relevant trait variation.

Here, we describe the development of fast neutron (FN) irradiated mutant lines in *Ipomoea purpurea*. First, we tracked germination, reproduction, and phenotypes across three generations of selfing following exposure to 5, 10, and 30 Gy of irradiation, relative to non-irradiated control replicates. Second, we conducted a mutant screen to assess the potential for visible mutant phenotypes, and determined if the phenotypic variance of traits related to the mating system, plant pollinator interactions and eco-physiology increased linearly with exposure to irradiation. We likewise identified outlier lineages, or those that exhibited extreme trait values relative to controls, and determined if fast neutron irradiation led to appreciable amounts of genetic variation among lines. Together, our results demonstrate that FN mutagenesis generates quantifiable variation in ecologically and evolutionarily relevant traits, providing a powerful resource for addressing questions in adaptation and agriculture.

## Methods

### Plant materials

To generate stable mutant lines using fast neutron (FN) irradiation, we first collected 60,000 seeds pooled across replicate plants of a single highly inbred (>20 generations) *Ipomoea purpurea* genotype (Line A) that was originally derived from a population collected from an agricultural field in Orange County, North Carolina, USA (Rausher and Fry 1993). High molecular weight DNA was isolated from the Line A genotype for PacBio HiFi sequencing and reference genome assembly.

We submitted seed to the University of California, Davis, McClellan Nuclear Research Center (McClellan, California, USA) for FN bombardment treatment at 5, 10, or 30 Gray (Gy). One Gray is equal to 1 Joule/kilogram of absorbed radiation dose. We then planted 500 seeds for each FN irradiation level (5, 10, or 30 Gy) to establish single-seed-descent genetic lines. These lines were propagated for three generations through self-fertilization in a growth room under a 7 hr daylength and 23°C to induce early flowering and rapid generation times (Fig. 1). This process decreases heterozygosity and increases the likelihood that phenotypes are heritable rather than due to physical damage from irradiation (Bolon et al. 2011). In addition, we propagated 500 lines through single-seed descent from untreated (0 Gy) seeds derived from the same Line A parent genotype. Across the irradiation treatments, we recorded the number of single-seed-descent lines that germinated and set seed each generation (Fig. 1). We likewise recorded potential mutant phenotypes each generation.

**Figure 1:**
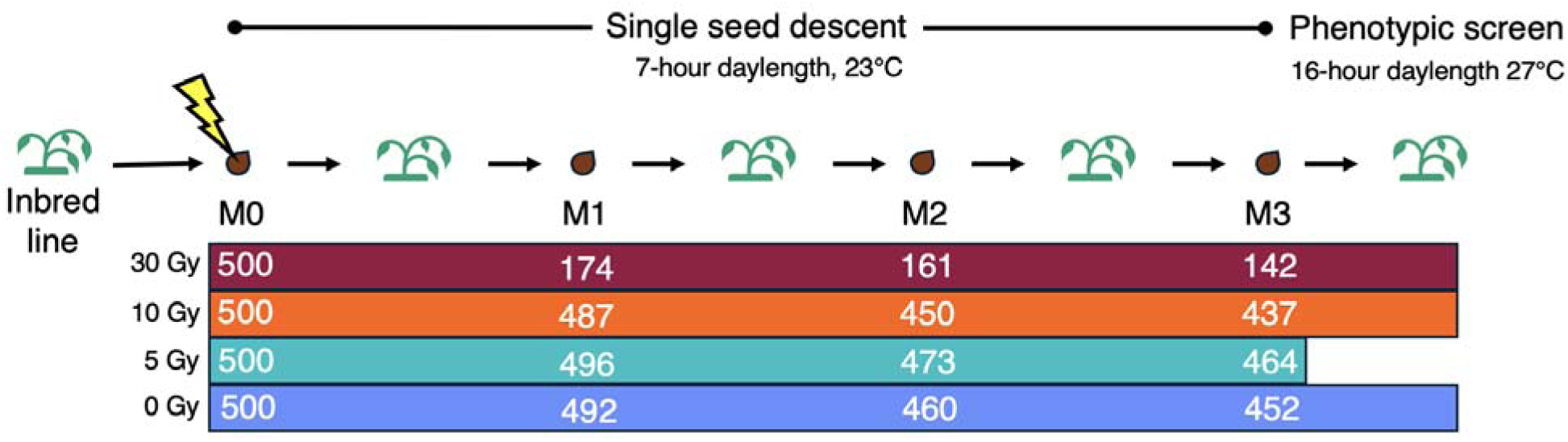
Diagram of the fast neutron line development of *Ipomoea purpurea*. Briefly, we collected thousands of seeds from a single inbred line. We then exposed 500 seeds to either 5, 10, or 30 Gy of fast neutron irradiation at the McClellan Nuclear Research Center (McClellan, California, USA) to establish individual lines. We planted these seeds as the M0 generation (original, irradiated seed) and proceeded with single-seed descent through three rounds of self-fertilization (M0, M1, and M2 were selfed) in the growth room under a 7-hour daylength and 23°C. Using the same method, we also established control lines (0 Gy) with three rounds of self-fertilization. To identify potential mutants and characterize phenotypic and genetic variation across irradiation treatment levels, we conducted a phenotypic screen of the M3 plants using the lines exposed to 0, 10, or 30 Gy under greenhouse conditions set to a 16-hour daylength and 27°C. We did not include lines exposed to 5 Gy in the phenotypic screen due to space constraints. Colored bars indicate the irradiation treatment level: 0 Gy (blue), 5 Gy (teal), 10 Gy (orange), and 30 Gy (dark red). The numbers within the bars reflect the number of lines at the start of that generation for each irradiation treatment level.

### Phenotypic screening and measurement across irradiation treatments

#### Experimental design and growth conditions

After three rounds of self-fertilization (M3), we phenotyped lines exposed to 0, 10, or 30 Gy to identify potential mutants and characterize phenotypic and genetic variation across irradiation treatment levels. We sowed three replicates of 100 lines from each treatment on October 14, 2024 under greenhouse conditions for a total of 900 individual plants. We re-sowed seeds on November 1 and 5, 2024 for 18 individuals that initially failed to germinate and we replaced 12 lines that had < 3 seeds with lines of the same irradiation dose. We excluded lines exposed to 5 Gy due to limited space.

We scarified and planted each seed in 10 cm pots, which were arranged in a completely randomized configuration across five benches in the greenhouse at Matthaei Botanical Gardens, University of Michigan (Ann Arbor, MI, USA). We grew the plants under supplemental lighting (16-hour days), with temperatures set to 27°C and bottom watering as necessary, from November 2024 to April 2025. To account for spatial variation in environmental conditions across the greenhouse, we divided each of the five benches into three or four quadrants using a 2×2 grid pattern, where each quadrant contained 36 to 48 plants.

Throughout the experiment, we examined each individual for visible mutant phenotypes (*i.e.*, plants with potentially abnormal phenotypes) and measured the following 14 floral and vegetative traits: date of first flowering, flower number, corolla width (mm) and length (mm), pollen count, pistil height (mm), height of tallest stamen (mm), leaf number, relative growth rate (leaves/day), leaf area (cm²), specific leaf area using dry mass (cm²/g), succulence using fresh mass (mg/cm²), trichome count, and anther-stigma distance (mm). Anther-stigma distance is calculated as height of the tallest stamen minus the height of the stigma, and is a trait associated with selfing rate in *Ipomoea purpurea* (Chang and Rausher 1998). Negative values reflect stigma length longer than the tallest anther resulting in an increased chance of outcrossing.

#### Floral and reproductive trait measurements

We calculated the date of first flowering as the first day an individual was recorded as flowering in reference to the planting day. Flower number represents the summed number of open flowers and buds that would open the next day, counted once weekly for 13 weeks. We took the floral size measurements (width and length of corollas, pistil height, stamen height) and assessed pollen production from 2,554 flowers from 718 individuals, such that at least two flowers were measured per individual on a given measurement day.

To assess pollen production, we collected anthers from each individual in separate microcentrifuge tubes, allowing the pollen to dehisce overnight with caps open. We then added 70% ethanol to preserve the pollen sample. Upon resuspending the pollen through mixing, we deposited 50 μl of the ethanol/pollen mixture onto a microscope slide, viewed the slide under a dissecting scope, and took an image using an iPhone camera. We used the multi-point tool in ImageJ (Schneider et al. 2012) to count the pollen grains. Therefore, our measure of pollen count reflects the number of pollen grains in a 50µl aliquot of the pollen sample.

#### Vegetative trait measurements: leaves and trichomes

We counted the number of leaves for each individual twice during the experiment, at ∼39 and ∼99 days post-sowing. We calculated the absolute growth rate (leaves/day) by subtracting the leaf count between the two censuses and dividing by the number of elapsed days (median 61 days) between the censuses. We evaluated the leaf number of each individual using the number of leaves counted ∼99 days post-sowing.

To assess leaf area, specific leaf area, succulence, and trichome count, we collected five mature leaves from each individual ∼112 days post-sowing. We placed the leaves in labeled bags, weighed them together to determine fresh mass, then scanned the leaves and placed them in a drying oven set to 60°C for at least 48 hours. At a later date, we weighed the dry leaves to determine dry mass. To calculate leaf area, we used the program Ilastik (Berg et al. 2019) and a custom Python script (Supplementary file 1). We calculated specific leaf area by dividing leaf area by dry leaf weight, and calculated succulence by subtracting dry leaf weight from fresh leaf weight and then dividing by leaf area (Reimann and Breckle 1995). We determined trichome count by imaging the adaxial surface of two leaves per individual using an iPhone mounted on a dissecting microscope (field of view = 0.2 cm^2^). Images were taken adjacent to the midrib, and non-glandular trichomes were counted using the multi-point tool in ImageJ (Schneider et al. 2012).

#### Screening and documentation of mutant phenotypes

During screens for potential mutants, we noted the number of replicates within a line that exhibited distinct mutant phenotypes to assess consistency within lines, and we photographed all plants with visible phenotypes. We documented leaf abnormalities using a flatbed scanner, and imaged a curled trichome mutant using scanning electron microscopy (TM3030, Hitachi High-Technologies Corporation, Tokyo, Japan). Specifically for line 1509, we grew additional plants from the same line (*i.e.*, full siblings) under a 7-hour daylength and 23°C in a growth room to obtain standardized images.

### Statistical analyses of phenotype measurements

For analysis, we averaged the values for corolla width and length, pollen count, stamen height, pistil height, anther-stigma distance, leaf number, leaf area, specific leaf area, succulence, and trichome count for each individual, since we took multiple measurements of these traits over the course of the experiment. Unless specified otherwise, all analyses were performed using R (ver. 4.5.2, R Core Team 2025) and RStudio (ver. 2026.07.0+139, Posit Team 2025), and results were visualized using ggplot2 (ver. 4.0.1, Wickham 2016).

### Phenotypic variance

We first assessed the amount of phenotypic variance per trait per treatment (0, 10, 30 Gy) to determine if irradiation increased variance according to exposure level. To do so, we estimated phenotypic variance for each trait within each irradiation treatment using a linear mixed model and bootstrapping approach.

We included bench quadrant (a combination of bench identity and quadrant position) and line as random effects in the models to account for spatial heterogeneity and genetic variation (lmer function of R package lme4, ver. 1.1-37,(Bates et al. 2015). We assessed model fit using the function simulateResiduals (R package DHARMa, ver. 0.4.7, (Hartig 2025). Each model followed this general structure:

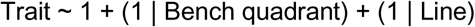

We extracted the residuals from each model (resid function of R package stats, ver. 4.5.2, R Core Team 2025), which represent trait variation after adjusting for block and line effects, and calculated the observed variance from these residuals. We then generated 10,000 bootstrap resamples of the residuals with replacement and calculated variance for each resample. We then constructed 95% confidence intervals for variance using the 2.5th and 97.5th percentiles of the bootstrap distribution.

### Detection of genetic variation and outlier identification

To identify potential genetic outliers for continuously distributed traits, we first analyzed each trait as a function of line with a covariate for block in separate linear mixed models for each irradiation treatment (lmer function of R package lme4). We assessed model fit using the function simulateResiduals (R package DHARMa). Each model followed this general structure:

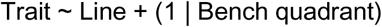

We determined the significance of Line using F tests with Satterthwaite’s approximation for degrees of freedom (R package lmerTest, ver. 3.1-3, Kuznetsova et al. 2017). A significant effect of line indicates the presence of genetic variation for a trait within an irradiation treatment. Because we assessed 14 traits, we applied a Benjamini-Hochberg false discovery rate correction to adjust the p-value (p.adjust function of R package stats). We then filtered for lines that had minimum and maximum trait values beyond one standard deviation of the mean trait value (*i.e.*, below mean − 1 SD or above mean + 1 SD) within the relevant irradiation treatment, to identify potential genetic outliers. Outlier lines were identified only for traits displaying significant genetic variation before correction (p < 0.05). This approach increases the likelihood that extreme phenotypes reflect heritable genetic differences induced by mutagenesis rather than environmental variation or measurement error.

## Results

### Reproductive success and line survival across generations of single seed descent

Reproductive success was monitored throughout generations and declined substantially at the highest irradiation level. Across generations, the number of lines retained to the M2 generation differed markedly among treatments. Relative to the 500 lines initially planted per treatment, 452 (90.4%) control lines, 464 (92.8%) 5 Gy lines, 437 (87.4%) 10 Gy lines, and 142 (28.4%) 30 Gy lines successfully produced seed through the M2 generation (Fig. 2, Supplementary Table 1). Below we provide a detailed description of germination, flowering, and seed set for the M0, M1 and M2 generations.

**Figure 2:**
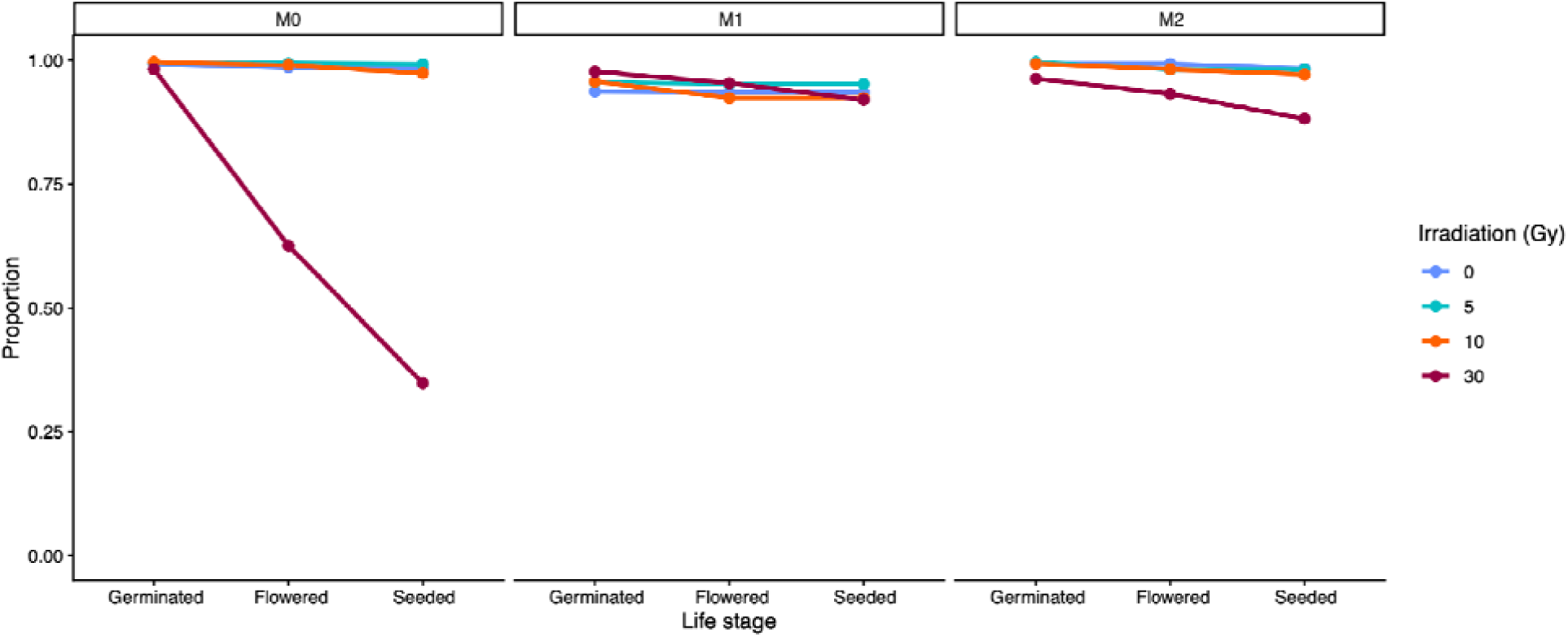
Visualization of line development under each irradiation treatment across the M0, M1 and M2 generations of single seed descent. Proportions are calculated with respect to the number of lines that were planted at the start of each generation (see Supplementary Table 1 for the corresponding numbers). Raw data points and connecting lines are plotted for the proportion of lines that germinated, flowered and set seed in the M0, M1 and M2 generations for the 0 Gy (blue), 5 Gy (teal), 10 Gy (orange), and 30 Gy (dark red) irradiation treatments. Ultimately, we generated M3 seed for a total of 1043 lines: 464 lines for 5 Gy, 437 for 10 Gy, and 142 for 30 Gy. In the phenotypic screen for the M3 generation, all of the lines that had at least one individual germinate, also flowered and set seed.

In the M0 generation, the 30 Gy lines showed reduced flowering and seed set. Germination was high across all treatments (0 Gy: 496, 99.2%; 5 Gy: 497, 99.4%: 10 Gy: 498, 99.6%; 30 Gy: 491, 98.2%). However, flowering was sharply reduced in the 30 Gy treatment, where 313 (62.6%) of the lines flowered compared to 493 (98.6%) in the control, 497 (99.4%) in 5 Gy and 495 (99.0%) in 10 Gy. Consequently, a further reduced 174 (34.8%) of the 500 originally planted 30 Gy lines set seed in the M0 generation compared to 492 (98.4%) in the control, 496 (99.2%) in 5Gy and 487 (97.4%) in 10 Gy. This reduced seed set was accompanied by a high frequency of floral abnormalities: 46.3% of flowering 30 Gy lines (145) exhibited abnormal flowers, compared to just 6.0% (30) and 3.8% (20) in the 5 Gy and 10 Gy treatments, respectively.

The pronounced decline in flowering and seed set observed for 30 Gy in M0 was absent in the M1 generation, though modest declines in line numbers persisted across all treatments. Because only lines that successfully set seed were advanced to subsequent generations, all generational percentages (M1 and M2) are calculated relative to the number of lines that reproduced in the previous generation.

Germination declined slightly in the M1 generation relative to M0, but remained high across treatments (0 Gy: 462, 93.9%; 5 Gy: 474, 95.5%; 10 Gy: 466, 95.7%; 30 Gy: 170, 97.7%). Flowering showed only a modest reduction compared to the M0 generation, with the lines in the 30 Gy treatment performing similar to other treatments (0 Gy: 460, 93.5%: 5 Gy: 472, 95.5%; 10 Gy: 466, 95.7%, 30 Gy: 166, 95.4%). The proportion of lines setting seed in the M1 generation remained high across treatments (0 Gy: 460, 93.5%: 5 Gy: 472, 95.2%; 10 Gy: 450, 92.4%, 30 Gy: 161, 92.5%)

Trends evident in the M2 generation mirrored those in the M1 generation, with consistent declines in lines advancing to the next life history stage across treatments. Germination remained high, exceeding 99% in the 0 Gy, 5 Gy, and 10 Gy treatments, with a modest reduction in the 30 Gy treatment (0 Gy: 457, 99.3%; 5 Gy: 471, 99.8%; 10 Gy: 447, 99.3%; 30 Gy: 155, 96.2%). Flowering proportions were similar to those observed in M1 (0 Gy: 457, 99.3%; 5 Gy: 464, 98.3%; 10 Gy: 447, 99.3%; 30 Gy: 150, 93.2%), as were seed set proportions (0 Gy: 452, 98.2%; 5 Gy: 464, 98.3%; 10 Gy: 437, 97.1%; 30 Gy: 142, 88.2%).

### Phenotypes observed during single seed descent (M0-M2 generations)

We report detailed descriptions of the phenotypes observed for each treatment in the M0-M2 generations in Supplementary file 2. Irradiation treatment induced diverse phenotypes across the single seed descent generations with radiation damage common in M0 lines, particularly at the 30 Gy treatment (see section: *Reproductive success and line survival across generations of single seed descent*). Most lines did not exhibit observable phenotypes by M1 and M2, suggesting recovery. Nevertheless, novel phenotypes continued to emerge in each generation in the 5 Gy and 10 Gy treatment levels.

### Visual phenotypes in the M3 screen

Our visual screening of the M3 generation cataloged distinct vegetative and floral mutant phenotypes but at a low frequency. Eleven mutant phenotypes (visually distinct relative to control lines) were observed within seven lines (3.5% of all the irradiated lines). We classified the phenotypes as vegetative and/or floral based on the affected developmental processes (Table 1, Fig. 3). Vegetative phenotypes encompassed three classes: developmental defects, pigmentation abnormalities and altered plant architecture. The developmental defects we observed were thin and small leaves, wrinkled leaves, curled trichomes, and small diameter stems. We also observed one pigmentation abnormality, where the plant looked to have variegated leaves, or the loss of chlorophyll in sections of the leaves. Plants with altered plant architecture displayed reduced leaf number, excessive leaf production and small size at flowering. We observed two classes of floral phenotypes, reproductive failure and floral defects. Lines with reproductive failure did not produce flowers, or had abnormal flowers that did not produce pollen. Lastly, one line with a floral defect produced functional flowers that were significantly smaller than wildtype (Fig. 3b).

**Figure 3:**
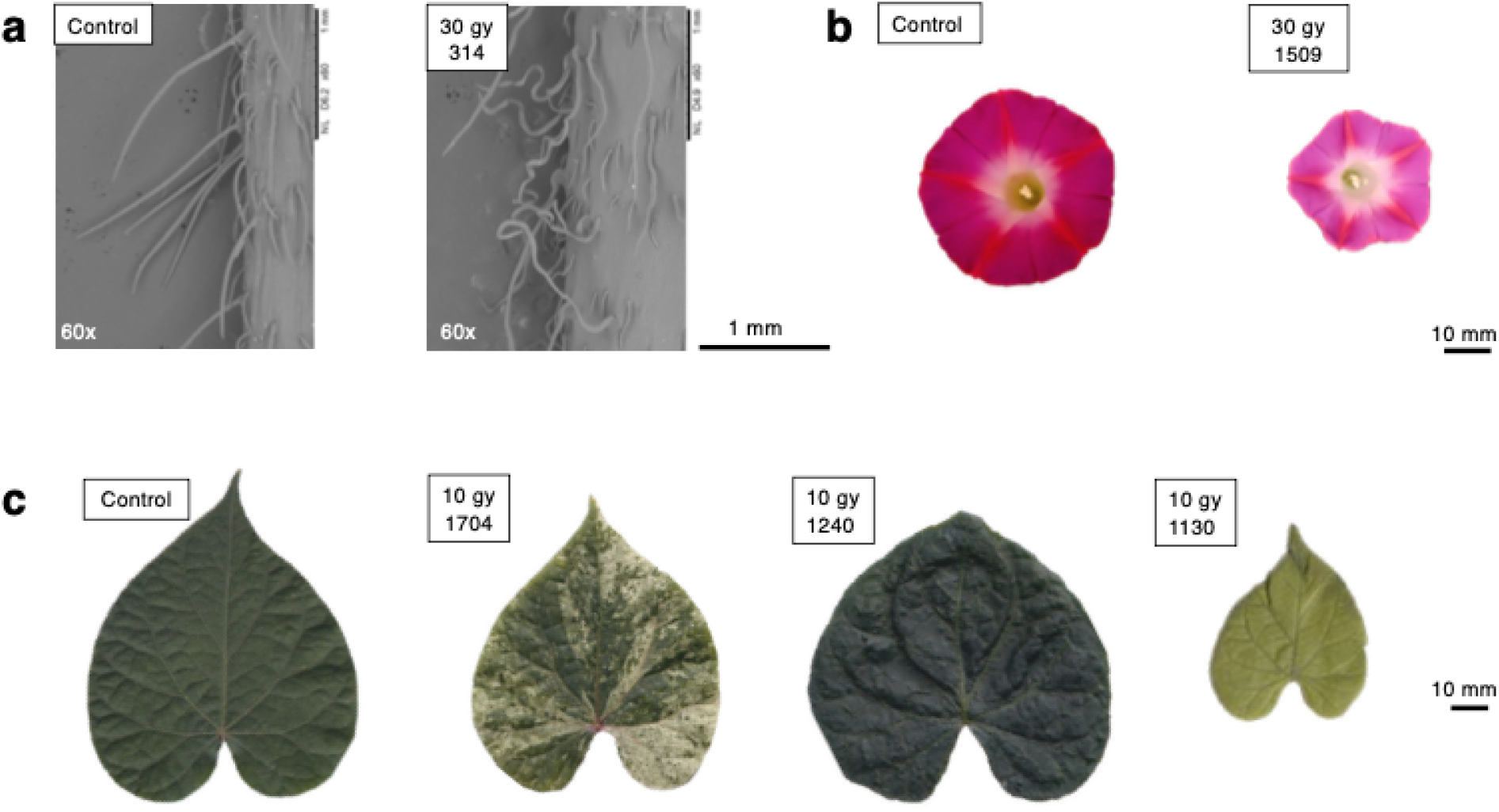
Images of select visual phenotypes observed in the M3 phenotype screen. a) scanning electron microscopy images in stem trichome morphology between a 0 Gy (control) line and 314, a line from the 30 Gy treatment with a developmental defect. b) Comparison of flower size and color between a control line and 1509, a line from the 30 Gy treatment with a floral defect. c) Comparisons of leaves between a control line and lines with developmental defects or pigmentation abnormalities in leaves: 1704 (10 Gy), 1240 (10 Gy), 1130 (10 Gy).

**Table 1:** Summary of visual phenotypes observed in the M3 phenotype screen. We observed 11 abnormal phenotypes across seven lines and classified the phenotypes as vegetative and/or floral based on the affected developmental processes. Lines 1130 and 1240 from the 10 Gy treatment, and lines 227 and 1509 from the 30 Gy treatment, displayed multiple visual phenotypes.

| Type | Description | Line | Gy | Seen in all individuals? |
| --- | --- | --- | --- | --- |
| Vegetative | Thin, small leaves | 1130 | 10 | No |
|  | Wrinkled leaves | 1240 | 10 | No |
|  | Variegated leaves | 1704 | 10 | No |
|  | Low number of leaves | 227 | 30 | Yes |
|  | Curling stem trichomes | 314 | 30 | Yes |
|  | Excessive leaf production and meristem growth | 1509 | 30 | Yes |
|  | Small diameter stem | 227 | 30 | Yes |
| Vegetative & floral | Small size at flowering | 1169 | 10 | Yes |
| Floral | No flower production | 1130 | 10 | No |
|  | Abnormal flowers that did not produce pollen | 1240 | 10 | No |
|  | Small flowers | 1509 | 30 | Yes |

Among the seven lines that we observed to have visual phenotypes, four exhibited consistent expression across individuals of up to three plants, while the remaining lines showed abnormalities for 1 or 2 individuals. Line 314 displayed curled trichomes on stems that were visible by eye, with normal flowers and leaves. Scanning electron microscopy revealed that these stem trichomes exhibited a ribbon-like morphology with twists. Comparatively, control plants produced straight trichomes (Fig. 3a). Line 1169 displayed a small size at flowering, where the individuals exhibited reduced stature and leaf production at the transition to flowering compared to controls. Line 1509 produced an excessive amount of leaves, coupled with lighter colored, smaller flowers (Fig. 3b). Additionally, these plants exhibited increased primary stem elongation that wrapped around the support stake multiple times while leaf morphology appeared normal.

### Phenotypic and genetic variation in quantitative traits

In addition to the induced qualitative variants described above for individual lines, FN radiation altered within-line phenotypic variance of quantitative trait values in a dose-dependent manner. While most measured traits exhibited unimodal distributions and individuals with extreme trait values present in all FN treatments (Fig. 4), phenotypic variance increased with irradiation treatment for five traits: pistil height, anther-stigma distance, pollen count, flower number and leaf number. Specifically, for anther-stigma distance, pollen count, and flower number, confidence intervals were not overlapping between the control (0 Gy) and the 30 Gy treatment (Fig. 5, Supplementary Table 2). Anther-stigma distance variance increased progressively from 0.452 in the control treatment (95% CI 0.356 - 0.554), to 0.566 in the 10 Gy treatment (95% CI 0.427 - 0.713), and 0.771 in the 30 Gy treatment (95% CI 0.572 - 0.998). Similarly, pollen count variance rose from 143.604 (95% CI 115.534-173.971) in controls to 186.078 (95% CI 151.014-227.030) at 10 Gy and 221.222 (95% CI 176.358-269.137) at 30 Gy. Flower number variance showed the same dose-dependent pattern, increasing from 23.601 (95% CI 19.007-28.683) in the control treatment to 33.452 (95% CI 27.998-39.067) at 10 Gy and 43.820 (95% CI 33.439-56.979) at 30 Gy.

**Figure 4:**
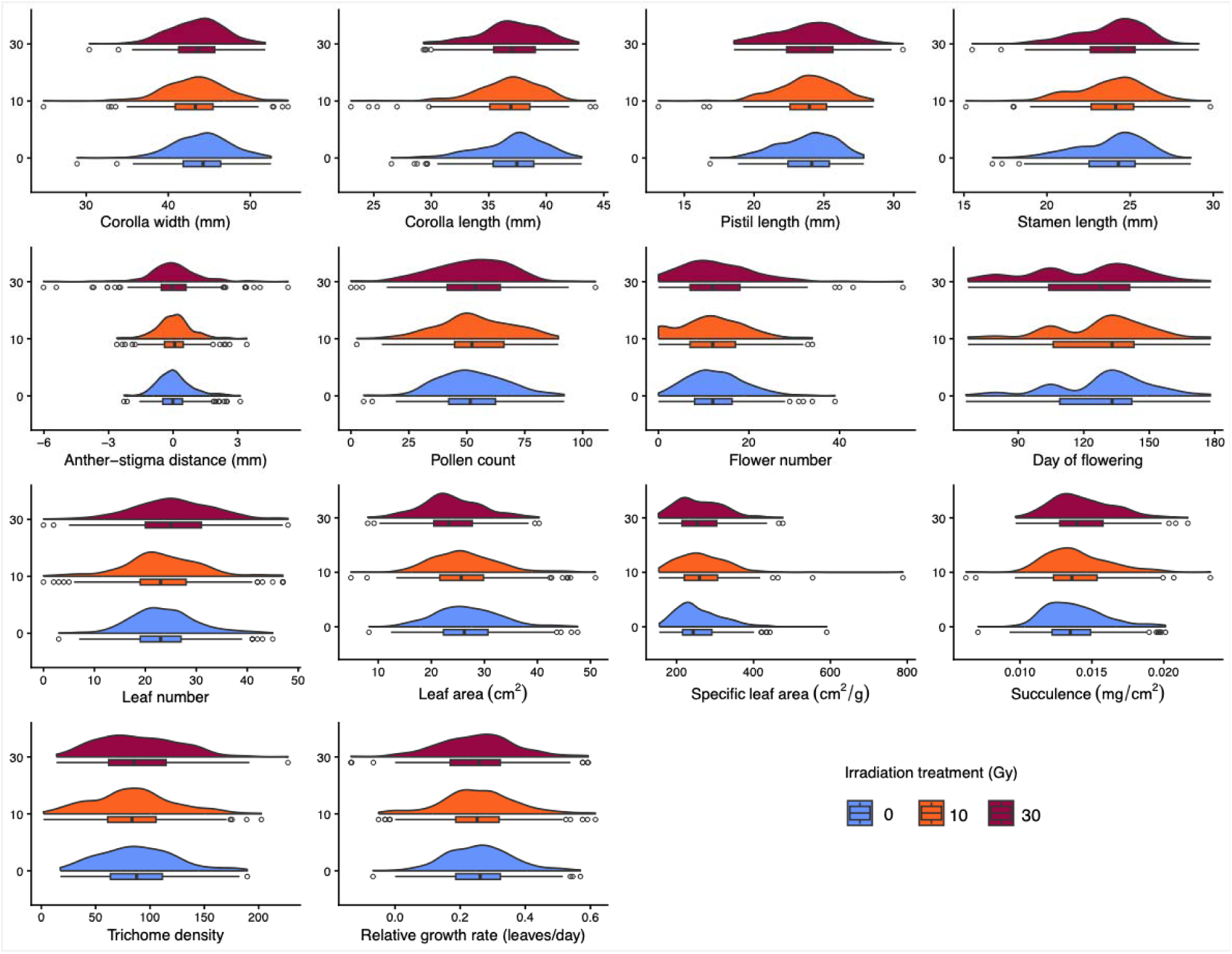
Distributions for each of the 14 traits measured in the M3 phenotypic screen. Violin plots and box plots display individual level data from 0 Gy (blue), 10 Gy (orange), and 30 Gy (dark red) irradiation treatments. White data points that reflect outlier measurements that fell more than 1.5 times the interquartile range (IQR) below the first quartile (Q1) or above the third quartile (Q3).

**Figure 5:**
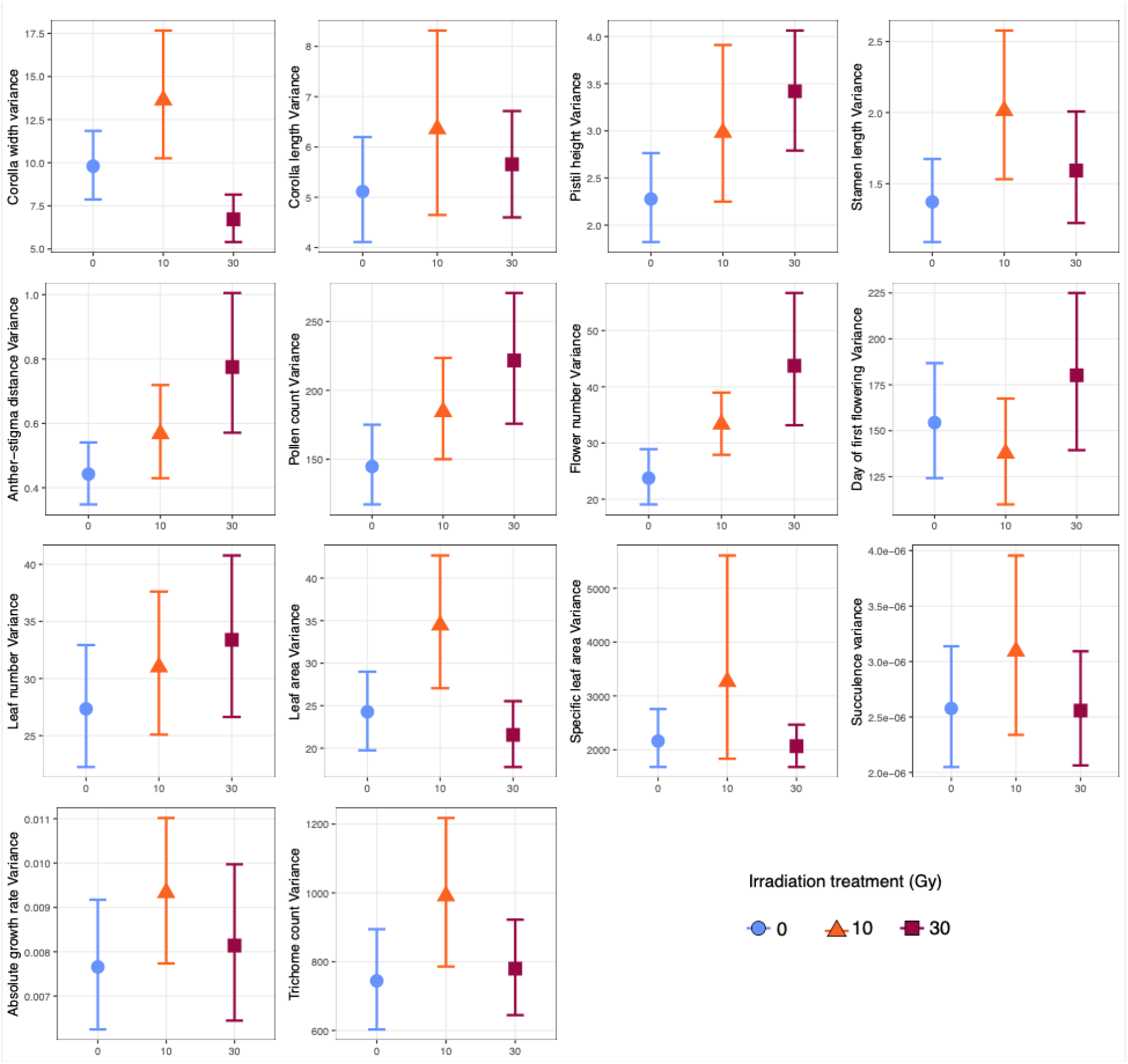
Phenotypic variance for each of the 14 continuous traits measured in the M3 phenotypic screen. Variance estimates are plotted for the 0 Gy (blue), 10 Gy (orange), and 30 Gy (dark red) irradiation treatments. Confidence intervals were constructed from the 2.5th and 97.5th percentiles of bootstrap distributions generated from 10,000 resamples with replacement of model residuals. Supplementary Table 2 contains the values for the variance and confidence intervals calculated for each trait and irradiation treatment.

Interestingly, the 30 Gy treatment reduced variance for corolla width and leaf area compared to 10 Gy treatment (Fig. 5). Corolla width variance was 9.781 (95% CI 7.891-11.867) in controls, increased to 13.729 (95% CI 10.398-17.789) at 10 Gy, but decreased to 6.734 (95% CI 5.382-8.176) at 30 Gy. Leaf area variance followed a similar pattern, measuring 27.172 (95% CI 22.057-32.540) in controls, increasing to 34.527 (95% CI 27.084-42.723) at 10 Gy, then declining to 21.782 (95% CI 17.995-25.738) at 30 Gy (Fig. 5, Supplementary Table 2).

If irradiation increased genetic variation for a trait, we would expect greater differentiation among lines within a treatment at higher doses. To test this, we treated Line as a fixed effect and assessed its significance using F-tests with Satterthwaite’s approximation for degrees of freedom. We detected significant genetic variation among lines, with highest radiation doses producing greater phenotypic differentiation between lines (Supplementary Table 3). The first day of flowering showed significant line effects across all treatments, including the control (0 Gy: F_99_ = 1.388, p = 0.030; 10 Gy: F_95_ = 1.751, p = 0.001; 30 Gy: F_98_ = 1.585, p = 0.005). The control treatment did not retain significance after p-value correction. The 10 Gy treatment had three additional traits with significant genetic variation prior to p-value correction (i.e. a significant line effect): flower number (F_99_ = 1.448, p = 0.016), leaf number (F_94_ = 1.488, p = 0.014), and succulence (F_95_ = 1.446, p = 0.022). In contrast, the 30 Gy treatment exhibited nine traits with significant levels of genetic variation: corolla width (F_98_ = 1.384, p = 0.045), flower number (F_99_ = 1.343, p = 0.044), pistil height (F_98_ = 1.529, p = 0.013), stamen length (F_98_ = 1.576, p = 0.008), Anther-stigma distance (F_98_ = 2.289, p < 0.001), leaf number (F_98_ = 1.725, p = 0.001), succulence (F_98_ = 1.382, p = 0.034), leaf area (F_98_ = 1.799, p < 0.001), and absolute growth rate (F_97_ = 1.446, p < 0.018).

After p-value correction for false discovery rate, the 10 Gy treatment retained significant line effects for only the first day of flowering (p = 0.016) and the 30 Gy treatment retained significant line effects for eight traits: day of flowering (p = 0.022), pistil height (p = 0.032), stamen length (p = 0.023), anther stigma distance (p < 0.001), absolute growth rate (p = 0.031), leaf area (p = 0.227), leaf number (p = 0.002), and succulence (p = 0.032).

### Identification of extreme and multi-trait outlier lines

We identified outlier lines as those deviating more than one standard deviation from the treatment mean for traits showing significant genetic variation prior to p-value correction (Table 2, See Supplementary Table 4 for line-level detail). Eight traits had outlier lines with varying frequencies: corolla width (4 lines, all below treatment mean), stamen length (5 lines, distributed below and above treatment mean), pistil height (6 lines, distributed both below and above treatment mean), anther-stigma distance (1 line, below treatment mean), flower number (4 lines, distributed below and above treatment mean), leaf number (4 lines, distributed below and above treatment mean), leaf area (3 lines, distributed below and above treatment mean), and succulence (7 lines, distributed below and above treatment mean). No outliers were identified for leaf number in the 10 Gy treatment. Outlier lines identified for all traits irrespective of genetic variation levels are presented in Supplementary Figure 1.

**Table 2.** Summary of outlier lines identified for traits with significant genetic variation within a treatment level. Outlier lines were defined as those whose line mean trait values fell more than one standard deviation above or below the treatment-level mean. “Direction” indicates whether outlier line means fell below, above, or on both sides of the treatment mean. Line-level identities, individual trait means, and standard deviations are reported in Supplementary Table 4.

| Trait | Gy | No. outlier lines | Direction | Range of line means |
| --- | --- | --- | --- | --- |
| Corolla width (mm) | 30 | 4 | All below mean | 36.8 – 39.6 |
| Pistil height (mm) | 30 | 6 | Below and above mean | 19.9 – 28.96 |
| Flower number | 10, 30 | 4 | Below and above mean | 0.34 – 25 |
| Leaf area (cm<sup>2</sup>) | 30 | 3 | Below and above mean | 12.8 – 36.3 |
| Stamen length (mm) | 30 | 5 | Below and above mean | 18.05 – 27.17 |
| Anther-stigma distance (mm) | 30 | 1 | Below mean | -4.84 |
| Leaf number | 30 | 4 | Below and above mean | 9.34 – 38 |
| Succulence (mg/cm<sup>2</sup>) | 10, 30 | 7 | Below and above mean | 11.4 – 17.4 |

Three lines from the 30 Gy treatment displayed outlier phenotypes across multiple quantitative traits (Supplementary Table 4). Line 1509 exhibited the most extensive multi-trait phenotype, with smaller flowers (mean 36.80 mm ± 1.65 SD; 30 Gy mean: 43.44 mm ± 3.37 SD), reduced stamen length (20.03 mm ± 0.99 SD; 30 Gy mean: 23.87 mm ± 2.06 SD), reduced pistil height (20.13 mm ± 1.12 SD; 30 Gy mean: 23.96 mm ± 2.41 SD), and increased leaf production (38.00 leaves ± 1.63 SD; 30 Gy mean: 25.42 leaves ± 7.98 SD). Line 1508 similarly had reduced stamen (20.44 mm ± 1.37 SD) and pistil height (19.9 mm ± 0.8 SD). Line 1499 showed the opposite pattern for stamen length and pistil height, exceeding the 30 Gy mean (stamen length: 27.17 mm ± 0.23 SE; pistil height: 28.43 mm ± 0.39 SD).

## Discussion

In this study, we document the first fast neutron mutant library for *Ipomoea purpurea.* We generated a total of 1043 M3 generation lines across 5 Gy, 10 Gy and 30 Gy irradiation doses. To characterize this resource, we pursued five aims: we tracked germination, reproduction, and phenotypes from the M0 through M2 generation to document line development across all four dose treatments; conducted a phenotypic screen of M3 lines to identify discrete visible mutants; tested whether fast neutron irradiation increases phenotypic and genetic variance across traits related to mating system, plant-pollinator interactions, and eco-physiology; and identified outlier lines exhibiting extreme trait values. By generating lines across a range of doses and applying a variance-based analytical framework, this resource adds to the utility of fast neutron mutagenesis and complements existing genomic approaches for linking mutations to trait variation and, ultimately, to consequences in natural populations.

### Line survival and phenotype recovery across generations

Line survival and phenotype expression across the M0–M2 generations were clearly dose-dependent, with the 30 Gy treatment producing substantially greater attrition than the 5 Gy and 10 Gy treatments. Line retention to the M3 generation declined with increasing dose, with the most severe attrition occurring at the M0 generation in the 30 Gy treatment (Fig. 2), mirroring patterns reported in fast neutron populations in *Glycine max* (Bolon et al. 2011). This attrition was accompanied by a high frequency of severe phenotypic disruption: 53.5% of flowering 30 Gy lines exhibited floral abnormalities in the M0 generation, compared to just 0.6% and 0.4% in the 5 Gy and 10 Gy treatments, respectively.

The phenotypes observed across the M0–M2 generations also revealed a temporal pattern with implications for how induced mutations become detectable during single seed descent. Radiation damage was common among M0 individuals, particularly at 30 Gy, but most of these early phenotypes were not recovered in subsequent generations. This is consistent with M0 plants representing chimeric tissue derived from irradiated seed in which physiological injury and non-heritable cellular damage are present along with induced mutations (Nielen et al. 2018). In contrast, novel phenotypes continued to appear in the 5 Gy and 10 Gy treatments through the M1 and M2 generations, well after this initial wave of injury had resolved. This delayed emergence is consistent with the expected dynamics of recessive mutation segregation under repeated self-fertilization: heterozygous mutations induced in the M0 generation are not phenotypically visible until they become homozygous, a state reached only after one or more generations of selfing. Because our phenotypic screen targeted the M3 generation, our results reflect the expression of recessive mutations rather than the transient physiological damage that is present in the initial M0 generation.

### Visual mutant phenotypes

In the visual screen of the M3 generation, we identified seven mutant lines exhibiting a total of 11 discrete visual phenotypes, indicating that some lines expressed more than one mutant trait (Table 1). These visually screened lines represented 3.5% of all irradiated lines, consistent with proportions reported in other fast neutron populations. For example, Bolon and colleagues documented visual phenotypes in approximately 2% of M2 lines in *Glycine max* (Bolon et al. 2011). This rate is also similar to those reported in other mutant screens, including T-DNA insertion lines in *Arabidopsis thaliana* (Wilson-Sánchez et al. 2014; Rutter et al. 2019). The phenotypic classes we observed were also broadly similar to those reported in other fast neutron screens, including defects in trichome development and growth defects in *G. max* (Bolon et al. 2011) and *Medicago truncatula* (Du et al. 2021). This overlap likely reflects two non-mutually exclusive factors: disruption of conserved developmental pathways that produce recognizable phenotypes across species, and an inherent detection bias in visual screens toward more striking morphological abnormalities. It is worth noting that visual screening identifies only the subset of mutant lines with discrete, scorable phenotypes, and that lines exhibiting subtler quantitative trait perturbations were identified through our variance-based analyses.

### Phenotypic and genetic variation in quantitative traits

Many traits tied to adaptation and crop improvement are polygenic, with phenotypic outcomes shaped by the additive contributions of many loci. We therefore examined whether fast neutron irradiation increased trait variance among mutant lines relative to non-irradiated controls across a suite of traits of interest to evolutionary biologists, including mating system traits, plant-pollinator interactions, and eco-physiological traits. Because the parental line used in this study was highly inbred prior to irradiation, standing genetic variation is negligible, allowing observed increases in trait variance to be attributed more directly to irradiation-induced mutations.

We found that phenotypic variance increased with irradiation across seven traits, with three, anther-stigma distance, pollen count, and flower number, displaying non-overlapping confidence intervals across treatments. Six of the seven traits, anther-stigma distance, pistil height, stamen length, corolla width, pollen count, and flower number, are directly connected to floral development and reproductive function with trichome density as the only exception. The concentration of variance responses in floral traits may reflect a shared underlying genetic architecture. Floral traits in Ipomoea purpurea are likely under related regulatory control given the coordinated development of the corolla, stamens, pistil, and pollen (Theissen and Melzer 2007). For example, MADS-box transcription factors are known to control multiple floral organ identities simultaneously across angiosperms (Theissen and Melzer 2007), such that a single mutation affecting such a locus could increase variance across multiple floral traits through pleiotropy (Smith 2016). Indeed, we did identify lines in the 30 Gy treatment with multiple outlier floral trait values. Line 1509 exhibited reductions across corolla width, stamen length, and pistil height, suggesting a mutation disrupting a shared floral regulatory locus. Line 1499 showed the opposite pattern for stamen length, and pistil height, further suggesting a shared regulatory architecture where mutations can perturb multiple floral traits in a coordinated manner regardless of direction. Indeed, while we have generally assumed that corolla width is a continuous trait in *Ipomoea purpurea*, it is also possible that, similar to *Mimulus* (Ding et al. 2017), a large-effect locus underlies corolla size variation in this species.

Our examination of trait distributions revealed that heterogeneous patterns of variance increase across traits. While most traits showed unimodal distributions with extreme values present across irradiated treatments, overall distributions varied by trait. Anther-stigma distance was notable in that the 30 Gy distribution extended in both directions relative to the control, meaning there were both individuals with stigmas taller than anthers indicative of negative values, or increased potential for outcrossing, and lines with more positive values, representing increased self-fertilization capacity. Comparatively, flower number and pollen count had right-skewed distributions with elevated frequencies of outliers in 5 and 10 Gy treatments. Notably, corolla width, leaf area, trichome density, and specific leaf area showed lower variance at 30 Gy than at 10 Gy, a pattern we attribute to the attritioned line numbers at the highest irradiation dose excluding the most severely affected phenotypes and deflating variance estimates.

To assess whether fast neutron irradiation introduced heritable genetic variation in *I. purpurea*, we tested for significant among-line differences in trait values. Prior to corrections for multiple tests, we detected the presence of genetic variation for ten traits, with eight remaining significant post-correction. This confirms that fast neutron irradiation can generate heritable variation that is consistent enough within lines to be detected, but to also have implications for evolvability. Traits with fast neutron induced heritable variation provide tractable material for identifying underlying genomic loci. Of the ten traits, corolla width, pistil height, stamen length, anther-stigma distance, and flower number are all reproductive related, further supporting the notion that these floral traits could be sharing underlying genetic architecture. The remaining traits, leaf area, leaf number, absolute growth rate, and succulence, are vegetative and reflect resource acquisition and water use strategy. The heritable variation in these traits may reflect the disruption of developmental pathways related to vegetative growth, potentially including growth regulating factors known to control leaf size and initiation (Omidbakhshfard et al. 2018). Sequencing the outlier lines that we identified for the traits exhibiting genetic variation would allow for the identification of the causal mutations.

Traits that showed no increase in variance under either framework may reflect genetic compensation, developmental buffering, or stochastic sampling effects rather than an absence of induced mutation (Rutter et al. 2017). Collectively, these results demonstrate that fast neutron mutagenesis can generate variation in continuously distributed traits, reinforcing its potential as an underutilized tool for dissecting the genetic architecture of complex phenotypes.

### The survival-variance trade-off

Among the lines that survived to M3, the 30 Gy treatment yielded the greatest downstream phenotypic and genetic variance (*see Phenotypic and genetic variation in quantitative traits, above*), indicating that surviving lines carried mutations perturbing quantitative traits without fatally disrupting reproduction. This tradeoff is reflected directly in line yield: whereas the 30 Gy treatment suffered the greatest attrition of any dose (*see Line survival and phenotype recovery across generations*, *above*), it also produced only 142 M3 lines, compared to 437 and 464 lines in the 10 Gy and 5 Gy treatments, respectively, which is roughly a third as many lines despite disproportionately higher induced variation. For researchers developing fast neutron resources, this tradeoff suggests that higher irradiation doses can be a productive strategy for maximizing induced genetic variation, provided the substantial line attrition associated with those doses is logistically and financially feasible.

### Future applications of this resource

The trait variance increases align with major research questions in ecology and evolution, suggesting several applications of this resource. For mating system evolution, we observed an increase in phenotypic variance and detected genetic variation for anther-stigma distance, which is a key determinant of self-fertilization rate (Chang and Rausher 1998), and for pistil height and stamen length. These FN lines offer new opportunities to study the fitness consequences of mating system variation, including inbreeding depression, and to map the genetic basis of floral morphology traits that mediate the transition between selfing and outcrossing. For plant-pollinator interactions, we found significant levels of genetic variation for corolla width and flower number, both of which influence pollinator attraction and resource allocation (Day Briggs and Anderson 2025), providing a foundation for studies of pollinator preference and the genetic architecture underlying floral display. García et al. (2023) demonstrated that water deficit alters both floral phenotypes and patterns of phenotypic selection in *I. purpurea*, with implications for how plant-pollinator interactions may evolve under climate change. Our lines with reduced corolla width and altered flower number could be included in water deficit and pollinator restriction experiments to test whether selection gradients connect to fitness differences.

The heritable variation in vegetative traits that we detected here also has direct utility for weed ecology research in *I. purpurea*. Drought significantly reduces photosynthetic efficiency, leaf number, and total leaf area in this species, with water-stressed plants showing roughly half the photosynthetic rate and leaf area of well-watered controls (Atala and Gianoli 2009). These reductions in resource acquisition are likely to limit competitive ability in agricultural environments under variable water availability. Identifying the genetic basis of performance under drought is directly relevant to predicting how this weed will respond to increasingly drought-prone conditions. By measuring performance of the FN lines under drought stress, researchers could identify mutations and begin to map the genomic regions underlying drought-relevant ecophysiological traits in this system.

Beyond these system-specific applications, our results demonstrate that fast neutron mutagenesis can generate heritable variation in continuous traits relevant to ecology and evolution. Therefore, this *I. purpurea* fast neutron library provides a platform for integrating mutagenesis approaches and mapping the genetic basis of adaptive traits into ecological and evolutionary research.

To facilitate broad use of this resource, we have created a publicly accessible website (https://sites.google.com/umich.edu/ipomoea-genomic-resources/) that contains all relevant information for the *I. purpurea* fast neutron mutant library. Researchers can request seed for lines from the M1 - M3 generations across the control and 5, 10 and 30 Gy irradiation treatments. The line numbers found in this study correspond to line IDs FN-Ipur-#### on the website. In addition, the website includes links to phenotypic data from the M3 greenhouse screen, and will contain data from future phenotyping efforts using the FN lines. Links to genomic resources are also available, including to the annotated Line A genome, and sequence information for select FN lines. This resource is part of a broader effort to develop genomic resources for *Ipomoea* species, which encompasses additional germplasm generation and genomic data across the genus. We encourage researchers working on questions in evolutionary ecology, plant-pollinator interactions, weed biology, and functional genomics to request seeds and engage with this resource.

### Conclusion

Fast neutron mutant populations are valuable resources for functional genomics in crop and model plant systems (Bolon et al. 2011), and our results demonstrate that this approach can be effectively extended to a weed and evolutionary ecology system. Here we show that fast neutron irradiation generates significant phenotypic and genetic variation in continuous traits relevant to ecology and evolution in I. purpurea, which establishes a key foundation for genotype-to-phenotype research in this system. This resource opens new experimental possibilities for longstanding questions in ecology and evolution, including the genetic basis of mating system variation, the evolution of plant-pollinator interactions, and the mechanisms underlying adaptation. More broadly, this work positions fast neutron mutagenesis as an underutilized but powerful tool for generating heritable variation in ecologically relevant traits, complementing existing genomic approaches and expanding the toolkit available to researchers working at the intersection of genetics, ecology, and evolution.

## Acknowledgements

This resource was developed with support from the U.S. National Science Foundation (NSF EDGE FGT award 2220214). Any opinions, findings, and conclusions or recommendations expressed here are those of the authors and do not necessarily reflect the views of the National Science Foundation.

## Data availability

Seeds and voucher material for the Line A parental genotype, and all M1, M2 and M3-generation fast neutron mutant lines described in this study, are available upon request from the corresponding author. Line availability, phenotypic data, and updates on ongoing genomic resource development for *Ipomoea purpurea* are maintained at https://sites.google.com/umich.edu/ipomoea-genomic-resources/.

The Line A reference genome assembly and associated PacBio HiFi sequencing reads generated in this study have been deposited at CoGe under Genome ID: 69912.

All phenotypic data and analysis code used to generate the results and figures in this manuscript are available at Dryad (DOI: 10.5061/dryad.vq83bk49b).

## Supplementary material described in this article is available as follows

Supplementary Figure 1: Box plots for each trait measured in the M3 phenotypic screen for all lines in the a) 10 Gy and b) 30 Gy treatments.

Supplementary Table 1: Summary of line development for control and fast neutron irradiated treatments.

Supplementary Table 2: Trait phenotypic variance and 95% confidence intervals across irradiation treatments.

Supplementary Table 3: Summary of analyses of genetic variance for traits measured in the M3 phenotypic screen.

Supplementary Table 4: Line-level detail for outlier lines identified for traits with significant genetic variation within a treatment level.

Supplementary File 1: Python script used to to analyze leaf area

Supplementary File 2: Descriptions of phenotypes observed in the M0, M1, and M2 generations.

## Author contributions

Conceptualization: RSB, JLM, WAP

Methodology: MIJ, RSB, JLM, WAP, MC

Formal analysis: MIJ (statistical analyses of phenotypic and genetic variance)

Investigation: MG (fast neutron line development and single seed descent); MIJ (greenhouse phenotyping, floral and vegetative trait measurement); MIJ (visual mutant screening and imaging, including scanning electron microscopy)

Resources: JLM (reference genome assembly, HMW DNA isolation); RSB (reference genome sequencing, greenhouse facilities, Matthaei Botanical Gardens)

Data curation: MIJ, RSB, JLM

Writing – original draft: MIJ, RSB, JLM, EAB, MC

Writing – review & editing: all authors

Visualization: MIJ, RSB, MC

Supervision: RSB, JLM

Funding acquisition: RSB, JLM, WAP

All authors read and approved the final manuscript.

## Supporting information

**Supplementary Figure 1:** Box plots for each trait measured in the M3 phenotypic screen for all lines in the a) 10 Gy and b) 30 Gy treatments.

**Supplementary Table 1:** Summary of line development for control and fast neutron irradiated treatments.

**Supplementary Table 2:** Trait phenotypic variance and 95% confidence intervals across irradiation treatments.

**Supplementary Table 3:** Summary of analyses of genetic variance for traits measured in the M3 phenotypic screen.

**Supplementary Table 4:** Line-level detail for outlier lines identified for traits with significant genetic variation within a treatment level.

**Supplementary Methods 1:** Python script used to analyze leaf area

**Supplementary Methods 2:** Descriptions of phenotypes observed in the M0, M1, and M2 generations.

**Supplementary Figure 1:**
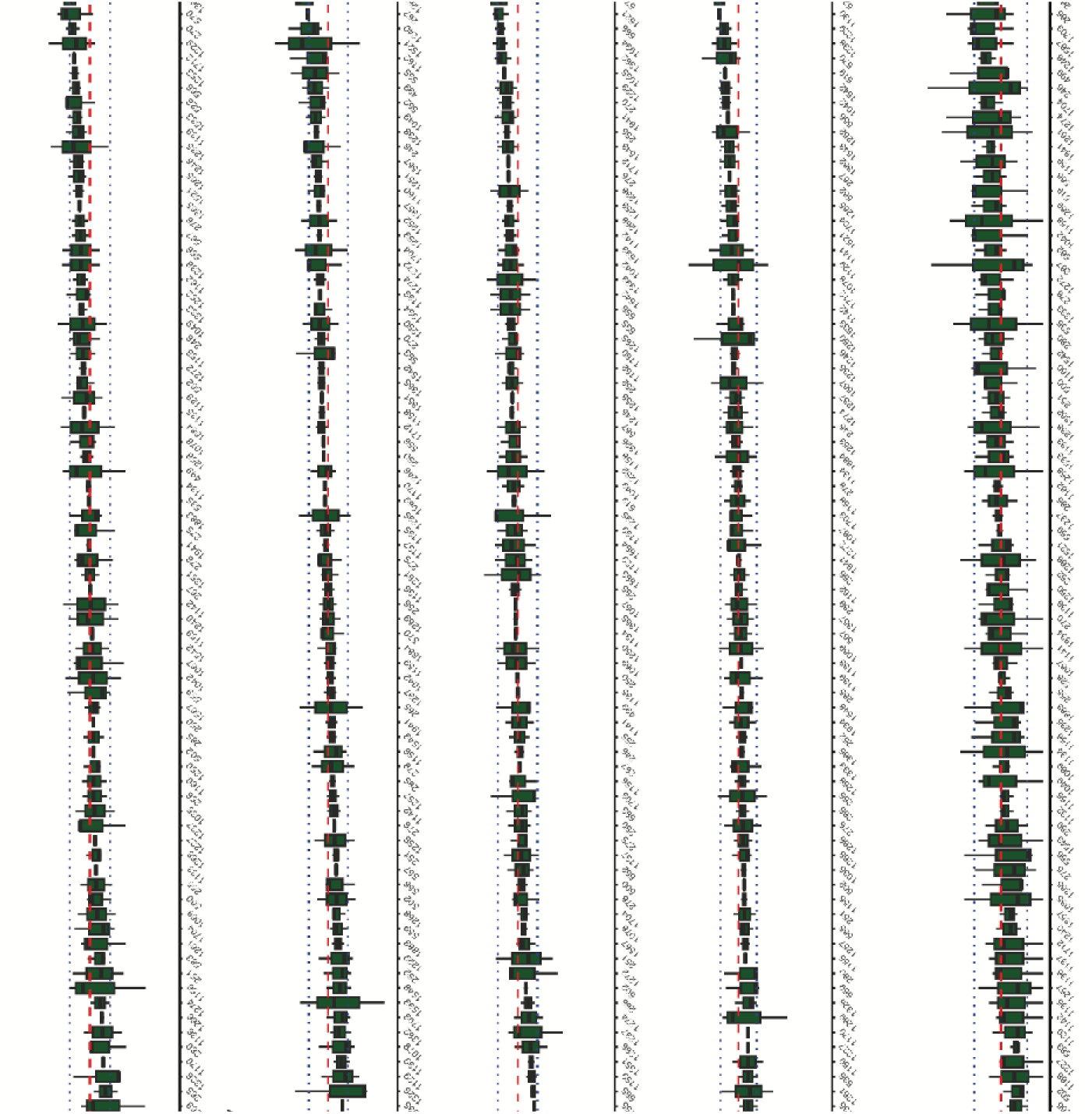
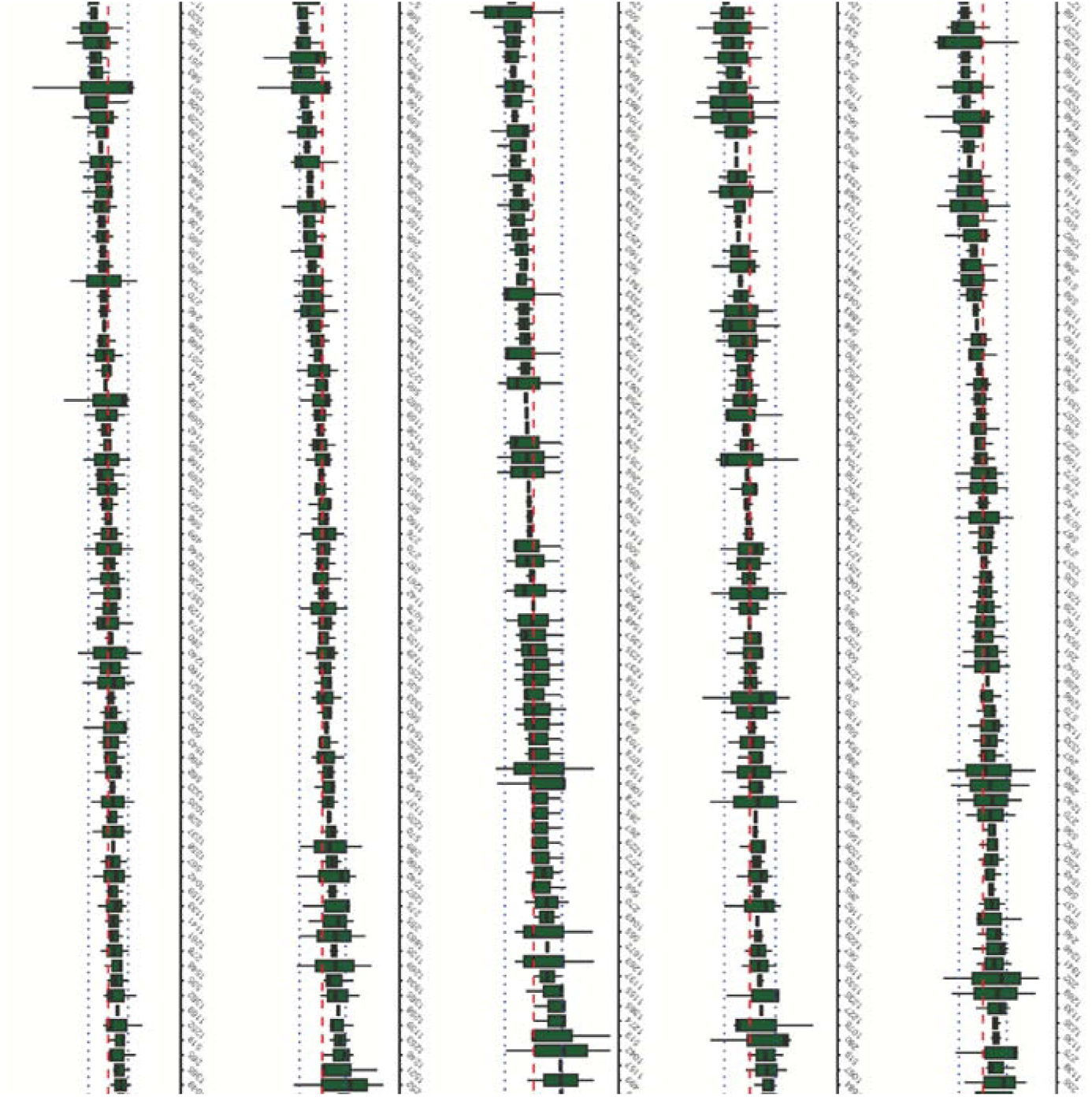
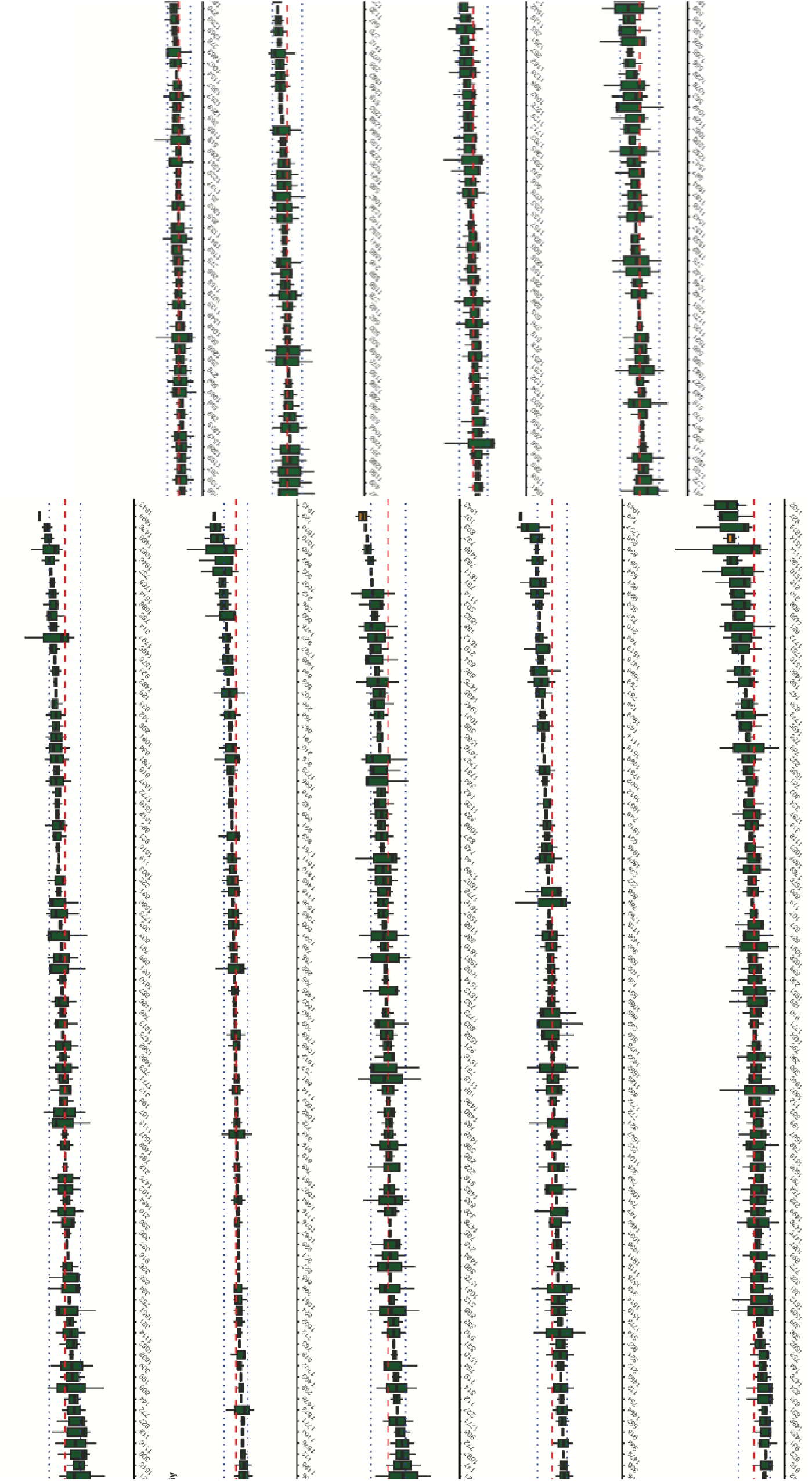
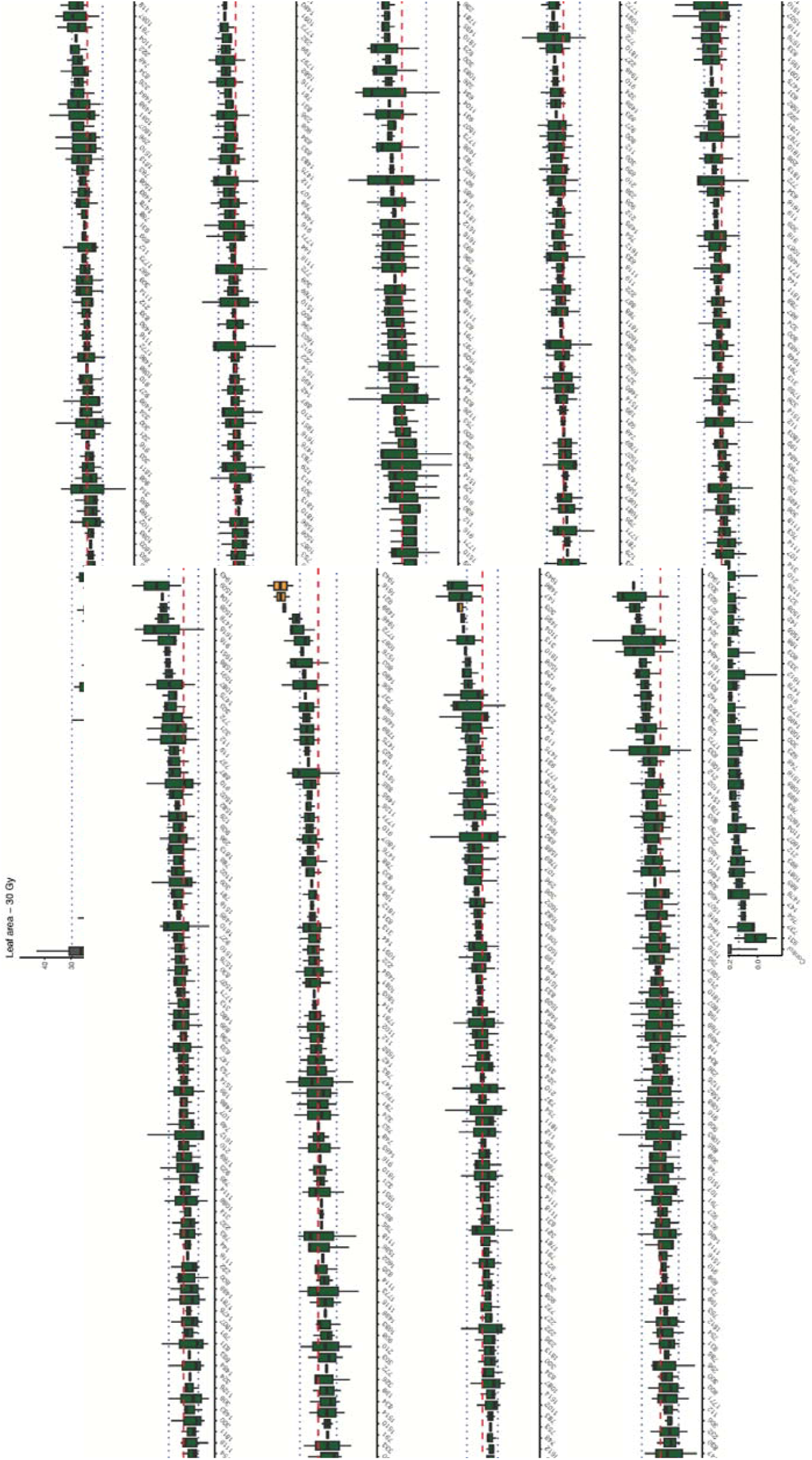
Box plots for each trait measured in the M3 phenotypic screen for all lines in the a) 10 Gy and b) 30 Gy treatments. Boxes display the mean value for each line with whiskers extending to the minimum and maximum values. Grey boxes represent control lines and are plotted first regardless of mean trait value. Yellow boxes indicate lines with trait values exceeding one standard deviation above or below the treatment mean. Green boxes indicate lines with trait values within one standard deviation of the treatment mean. The red line represents the treatment mean, and blue dotted lines represent one standard deviation above and below the treatment mean.

**Supplementary Table 1:** Summary of line development for control and fast neutron irradiated treatments. Initially, we started 500 lines for both the control lines and the fast neutron irradiated lines for single seed descent for a total of three generations. Lines declined per generation across treatments. Notably, 142, or 28.4%, of the initial 500 lines exposed to the 30 Gy treatment advanced to the M3 generation.

|  | Control |  |  | 5 Gy |  |  | 10 Gy |  |  | 30 Gy |  |  |
| --- | --- | --- | --- | --- | --- | --- | --- | --- | --- | --- | --- | --- |
| Generation | 0 | 1 | 2 | M0 | M1 | M2 | M0 | M1 | M2 | M0 | M1 | M2 |
| Lines planted | 500 | 492 | 460 | 500 | 496 | 473 | 500 | 487 | 450 | 500 | 174 | 161 |
| Lines germinated | 496 | 461 | 457 | 497 | 474 | 471 | 498 | 466 | 447 | 491 | 170 | 155 |
| Lines flowered | 493 | 460 | 457 | 497 | 472 | 464 | 495 | 450 | 442 | 313 | 166 | 150 |
| Lines seeded | 492 | 460 | 452 | 496 | 472 | 464 | 487 | 450 | 437 | 174 | 160 | 142 |

**Supplementary Table 2:**
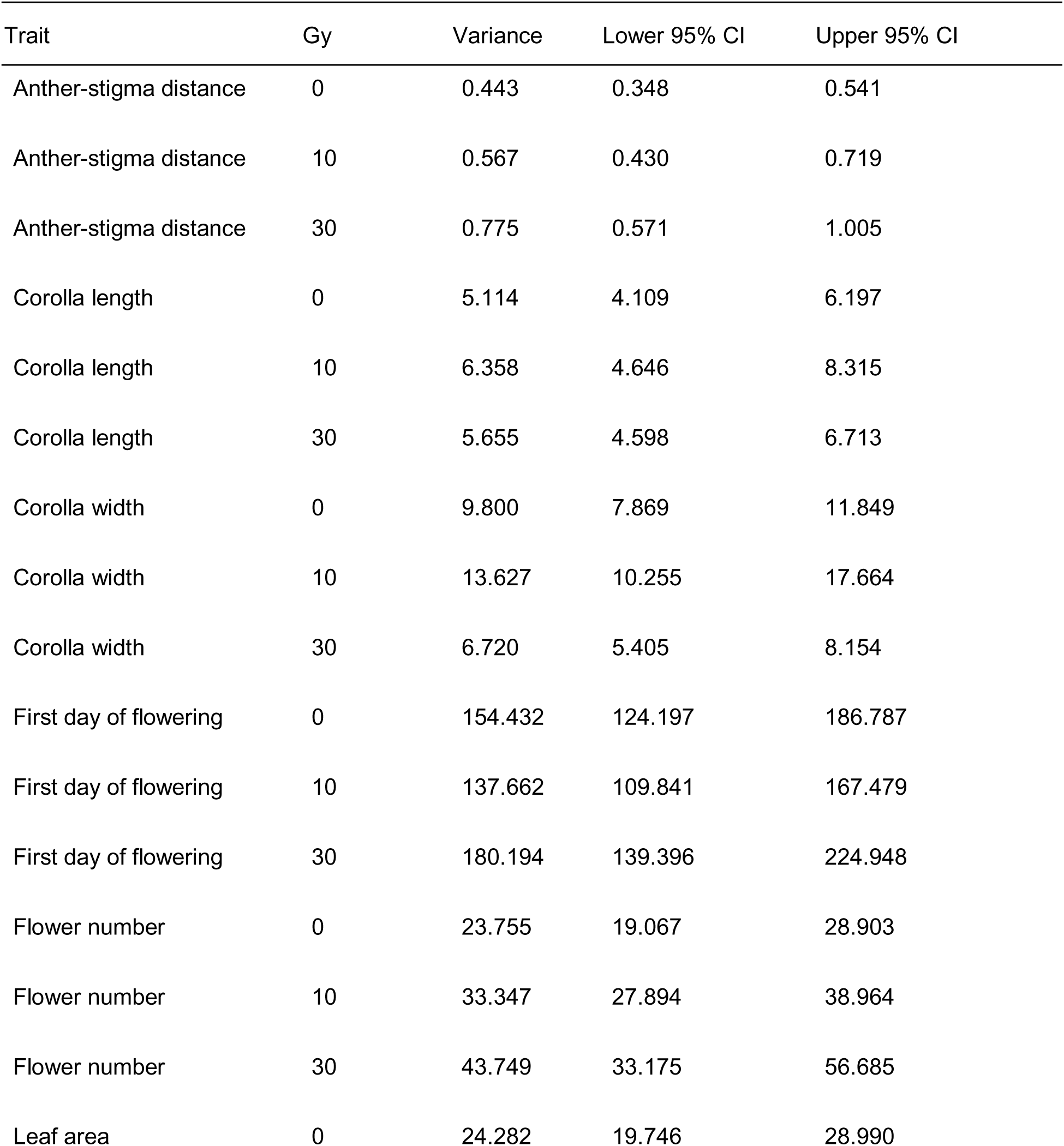
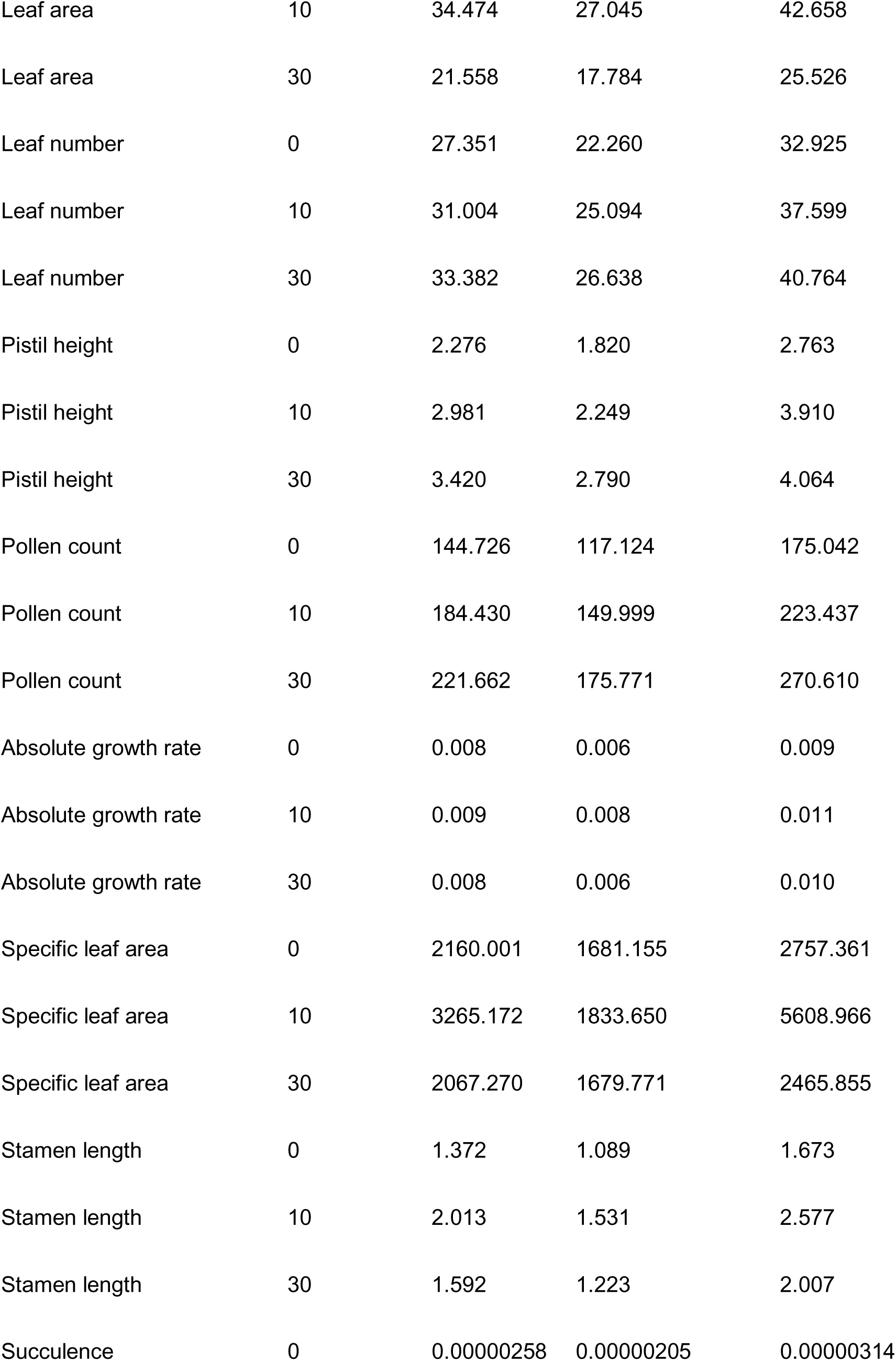
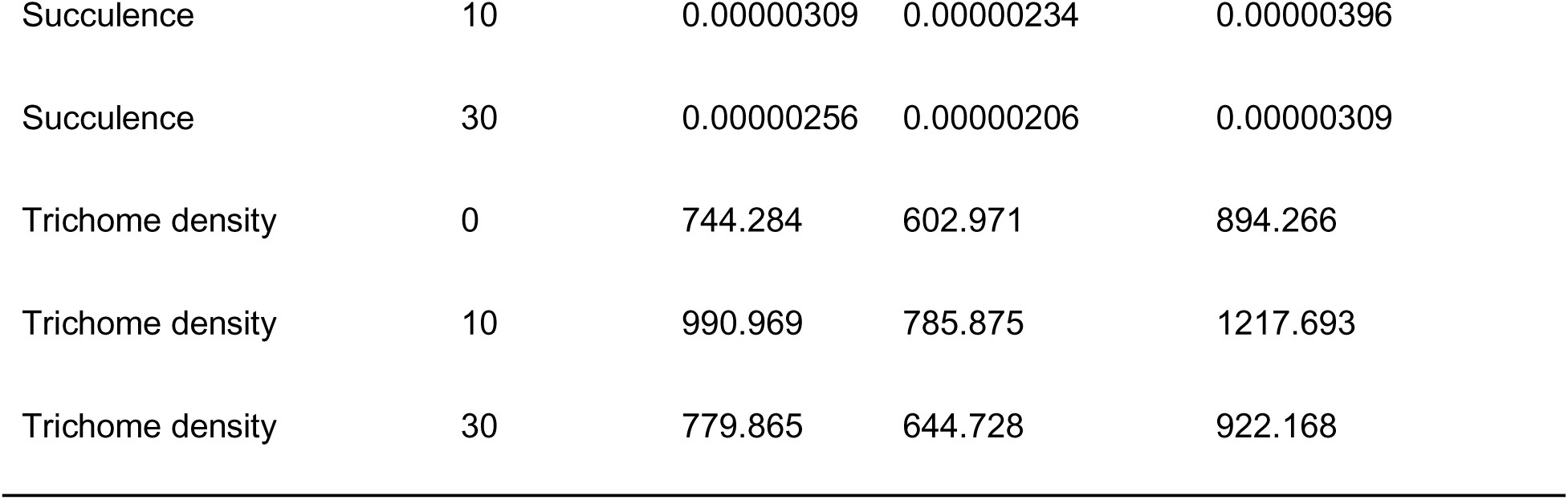
Trait phenotypic variance and 95% confidence intervals across irradiation treatments. We estimated phenotypic variance for each trait within each irradiation treatment using a linear mixed model and bootstrapping approach and included bench quadrant and line as covariates in the models to account for spatial heterogeneity and genetic variation (lmer function of R package lme4, Bates et al. 2015). Confidence intervals were constructed from the 2.5th and 97.5th percentiles of bootstrap distributions generated from 10,000 resamples with replacement of model residuals

| Trait | Gy | Variance | Lower 95% CI | Upper 95% CI |
| --- | --- | --- | --- | --- |
| Anther-stigma distance | 0 | 0.443 | 0.348 | 0.541 |
| Anther-stigma distance | 10 | 0.567 | 0.430 | 0.719 |
| Anther-stigma distance | 30 | 0.775 | 0.571 | 1.005 |
| Corolla length | 0 | 5.114 | 4.109 | 6.197 |
| Corolla length | 10 | 6.358 | 4.646 | 8.315 |
| Corolla length | 30 | 5.655 | 4.598 | 6.713 |
| Corolla width | 0 | 9.800 | 7.869 | 11.849 |
| Corolla width | 10 | 13.627 | 10.255 | 17.664 |
| Corolla width | 30 | 6.720 | 5.405 | 8.154 |
| First day of flowering | 0 | 154.432 | 124.197 | 186.787 |
| First day of flowering | 10 | 137.662 | 109.841 | 167.479 |
| First day of flowering | 30 | 180.194 | 139.396 | 224.948 |
| Flower number | 0 | 23.755 | 19.067 | 28.903 |
| Flower number | 10 | 33.347 | 27.894 | 38.964 |
| Flower number | 30 | 43.749 | 33.175 | 56.685 |
| Leaf area | 0 | 24.282 | 19.746 | 28.990 |
| Leaf area | 10 | 34.474 | 27.045 | 42.658 |
| Leaf area | 30 | 21.558 | 17.784 | 25.526 |
| Leaf number | 0 | 27.351 | 22.260 | 32.925 |
| Leaf number | 10 | 31.004 | 25.094 | 37.599 |
| Leaf number | 30 | 33.382 | 26.638 | 40.764 |
| Pistil height | 0 | 2.276 | 1.820 | 2.763 |
| Pistil height | 10 | 2.981 | 2.249 | 3.910 |
| Pistil height | 30 | 3.420 | 2.790 | 4.064 |
| Pollen count | 0 | 144.726 | 117.124 | 175.042 |
| Pollen count | 10 | 184.430 | 149.999 | 223.437 |
| Pollen count | 30 | 221.662 | 175.771 | 270.610 |
| Absolute growth rate | 0 | 0.008 | 0.006 | 0.009 |
| Absolute growth rate | 10 | 0.009 | 0.008 | 0.011 |
| Absolute growth rate | 30 | 0.008 | 0.006 | 0.010 |
| Specific leaf area | 0 | 2160.001 | 1681.155 | 2757.361 |
| Specific leaf area | 10 | 3265.172 | 1833.650 | 5608.966 |
| Specific leaf area | 30 | 2067.270 | 1679.771 | 2465.855 |
| Stamen length | 0 | 1.372 | 1.089 | 1.673 |
| Stamen length | 10 | 2.013 | 1.531 | 2.577 |
| Stamen length | 30 | 1.592 | 1.223 | 2.007 |
| Succulence | 0 | 0.00000258 | 0.00000205 | 0.00000314 |
| Succulence | 10 | 0.00000309 | 0.00000234 | 0.00000396 |
| Succulence | 30 | 0.00000256 | 0.00000206 | 0.00000309 |
| Trichome density | 0 | 744.284 | 602.971 | 894.266 |
| Trichome density | 10 | 990.969 | 785.875 | 1217.693 |
| Trichome density | 30 | 779.865 | 644.728 | 922.168 |

**Supplementary Table 3:** Summary of analyses of genetic variance for the 14 continuous traits measured in the M3 phenotypic screen. We analyzed each trait as a function of line with a covariate for block in separate linear mixed models for each irradiation treatment (lmer function of R package lme4). We determined the significance of Line using F tests with Satterthwaite’s approximation for degrees of freedom Because we assessed 14 traits, we also report the Benjamini-Hochberg false discovery rate corrected p value (P-FDR).

| Trait | Gy | N | F | P | P-FDR |
| --- | --- | --- | --- | --- | --- |
| Anther-Stigma distance | 0 | 258 | 1.311 | 0.070 | 0.336 |
| Anther-Stigma distance | 10 | 229 | 1.095 | 0.320 | 0.467 |
| Anther-Stigma distance | 30 | 237 | 2.291 | **0.00001** | **0.0001** |
| Corolla length | 0 | 257 | 0.900 | 0.710 | 0.792 |
| Corolla length | 10 | 227 | 1.218 | 0.155 | 0.363 |
| Corolla length | 30 | 234 | 1.207 | 0.163 | 0.175 |
| Corolla width | 0 | 257 | 1.032 | 0.427 | 0.552 |
| Corolla width | 10 | 227 | 1.066 | 0.368 | 0.467 |
| Corolla width | 30 | 234 | 1.393 | **0.0421** | 0.062 |
| First day of flowering | 0 | 290 | 1.388 | **0.0301** | 0.336 |
| First day of flowering | 10 | 260 | 1.751 | **0.0011** | **0.0155** |
| First day of flowering | 30 | 278 | 1.585 | **0.0048** | **0.0223** |
| Flower number | 0 | 300 | 1.026 | 0.434 | 0.552 |
| Flower number | 10 | 300 | 1.448 | **0.0158** | 0.074 |
| Flower number | 30 | 300 | 1.341 | **0.0443** | 0.062 |
| Leaf area | 0 | 295 | 1.288 | 0.072 | 0.336 |
| Leaf area | 10 | 259 | 1.050 | 0.390 | 0.467 |
| Leaf area | 30 | 280 | 1.529 | **0.0081** | **0.0227** |
| Leaf number | 0 | 299 | 1.140 | 0.222 | 0.442 |
| Leaf number | 10 | 265 | 1.488 | **0.0143** | 0.074 |
| Leaf number | 30 | 288 | 1.838 | **0.0002** | **0.0016** |
| Pistil height | 0 | 258 | 1.125 | 0.258 | 0.442 |
| Pistil height | 10 | 229 | 1.135 | 0.255 | 0.467 |
| Pistil height | 30 | 237 | 1.510 | **0.0150** | **0.0315** |
| Pollen count | 0 | 259 | 1.172 | 0.190 | 0.442 |
| Pollen count | 10 | 228 | 1.064 | 0.371 | 0.467 |
| Pollen count | 30 | 236 | 0.844 | 0.809 | 0.809 |
| Absolute growth rate | 0 | 299 | 0.891 | 0.736 | 0.792 |
| Absolutegrowth rate | 10 | 264 | 0.978 | 0.541 | 0.583 |
| Absolutegrowth rate | 30 | 284 | 1.446 | **0.0180** | **0.0315** |
| Specific leaf area | 0 | 295 | 0.789 | 0.903 | 0.903 |
| Specific leaf area | 10 | 259 | 0.942 | 0.619 | 0.619 |
| Specific leaf area | 30 | 280 | 1.191 | 0.161 | 0.175 |
| Stamen length | 0 | 258 | 1.225 | 0.133 | 0.442 |
| Stamen length | 10 | 229 | 1.314 | 0.080 | 0.225 |
| Stamen length | 30 | 237 | 1.602 | **0.0068** | **0.0227** |
| Succulence | 0 | 295 | 1.103 | 0.284 | 0.442 |
| Succulence | 10 | 259 | 1.446 | **0.0217** | 0.076 |
| Succulence | 30 | 280 | 1.451 | **0.0176** | **0.0315** |
| Trichome density | 0 | 294 | 1.111 | 0.271 | 0.442 |
| Trichome density | 10 | 258 | 1.046 | 0.401 | 0.467 |
| Trichome density | 30 | 280 | 1.251 | 0.103 | 0.131 |

**Supplementary Table 4:** Line-level detail for outlier lines identified for traits with significant genetic variation within a treatment level. Outlier lines were defined as those whose line mean trait values fell more than one standard deviation above or below the treatment-level mean. Population means (italicized) are shown for each irradiation treatment (0, 10, or 30 Gy) in which at least one outlier line was identified for that trait.

| Trait | Line | Gy | Trait mean ± SD |
| --- | --- | --- | --- |
| **Corolla width** (mm) | *Population mean* | 0 | 44.14 ± 3.52 |
| | *Population mean* | 30 | 43.44 ± 3.37 |
|  | 1087 | 30 | **39.6 ± 0.41** |
|  | 1616 | 30 | **39.5 ± 0.05** |
|  | 908 | 30 | **38.1 ± 1.3** |
|  | 1509 | 30 | **36.8 ± 1.65** |
| **Pistil height** (mm) | *Population mean* | 0 | 23.82 ± 2.04 |
| | *Population mean* | 30 | 23.96 ± 2.41 |
|  | 1508 | 30 | **19.9 ± 0.8** |
|  | 1509 | 30 | **20.13 ± 1.12** |
|  | 300 | 30 | **20.68 ± 0.47** |
|  | 1499 | 30 | **28.43 ± 0.39** |
|  | 921 | 30 | **28.83 ± 0.62** |
|  | 1616 | 30 | **28.96 ± 1.4** |
| **Flower number** | *Population mean* | 0 | 12.82 ± 6.4 |
| | *Population mean* | 10 | 11.68 ± 7.41 |
|  | 1169 | 10 | **0.34 ± 0.57** |
|  | 1347 | 10 | **0.34 ± 0.57** |
|  | 1672 | 10 | **0.67 ± 0.57** |
| | *Population mean* | 30 | 12.94 ± 8.26 |
|  | 1514 | 30 | **25 ± 3.61** |
| **Leaf area** (cm<sup>2</sup>) | *Population mean* | 0 | 26.75 ± 6.23 |
| | *Population mean* | 30 | 23.83 ± 5.81 |
|  | 227 | 30 | **12.8 ± 2.52** |
|  | 795 | 30 | **14.6 ± 0.67** |
|  | 1951 | 30 | **36.3 ± 4.19** |
| **Stamen length** (mm) | *Population mean* | 0 | 23.85 ± 2.08 |
| | *Population mean* | 30 | 23.87 ± 2.06 |
|  | 147 | 30 | **18.05 ± 3.03** |
|  | 1509 | 30 | **20.03 ± 0.99** |
|  | 1508 | 30 | **20.44 ± 1.37** |
|  | 908 | 30 | **20.66 ± 1.23** |
|  | 1499 | 30 | **27.17 ± 0.23** |
| **Anther-stigma distance (mm)** | *Population mean* | 0 | 0.05 ± 0.8 |
| | *Population mean* | 30 | 0.01 ± 1.31 |
|  | 147 | 30 | **-4.84 ± 1.57** |
| **Leaf number** | *Population mean* | 0 | 23.66 ± 6.54 |
| | *Population mean* | 30 | 25.43 ± 7.98 |
|  | 227 | 30 | **9.34 ± 2.08** |
|  | 1807 | 30 | **13 ± 2.64** |
|  | 754 | 30 | **14.67 ± 1.15** |
|  | 1509 | 30 | **38 ± 2.82** |
| **Succulence (mg/cm<sup>2</sup>)** | *Population mean* | 0 | 13.8 ± 2.09 |
| | *Population mean* | 10 | 14 ± 2.31 |
|  | 1237 | 10 | **11.4 ± 0.24** |
|  | 251 | 10 | **16.7 ± 0.76** |
| | *Population mean* | 30 | 14.3 ± 2.16 |
|  | 142 | 30 | **11.6 ± 0.69** |
|  | 1803 | 30 | **11.7 ± 0.29** |
|  | 893 | 30 | **11.7 ± 0.63** |
|  | 1495 | 30 | **17 ± 0.45** |
|  | 303 | 30 | **17.4 ± 0.71** |

**Supplementary File 1**: Python script used to analyze leaf area. Once images are processed in *ilastik* to distinguish leaves from background (and envelopes if needed), this script will count the pixels associated with the leaves and convert that to cm^2^ according to the scanner DPI, which should be 300 dpi.

This following command runs the script: python countLeafPixels_v3.py [argument1] [argument2]

argument1 the already-existing name of the lower-level / sub-directory (folder) containing all the image files (.tiff) that you want to process

argument2 a new name of the output file you want to generate with the results of the script

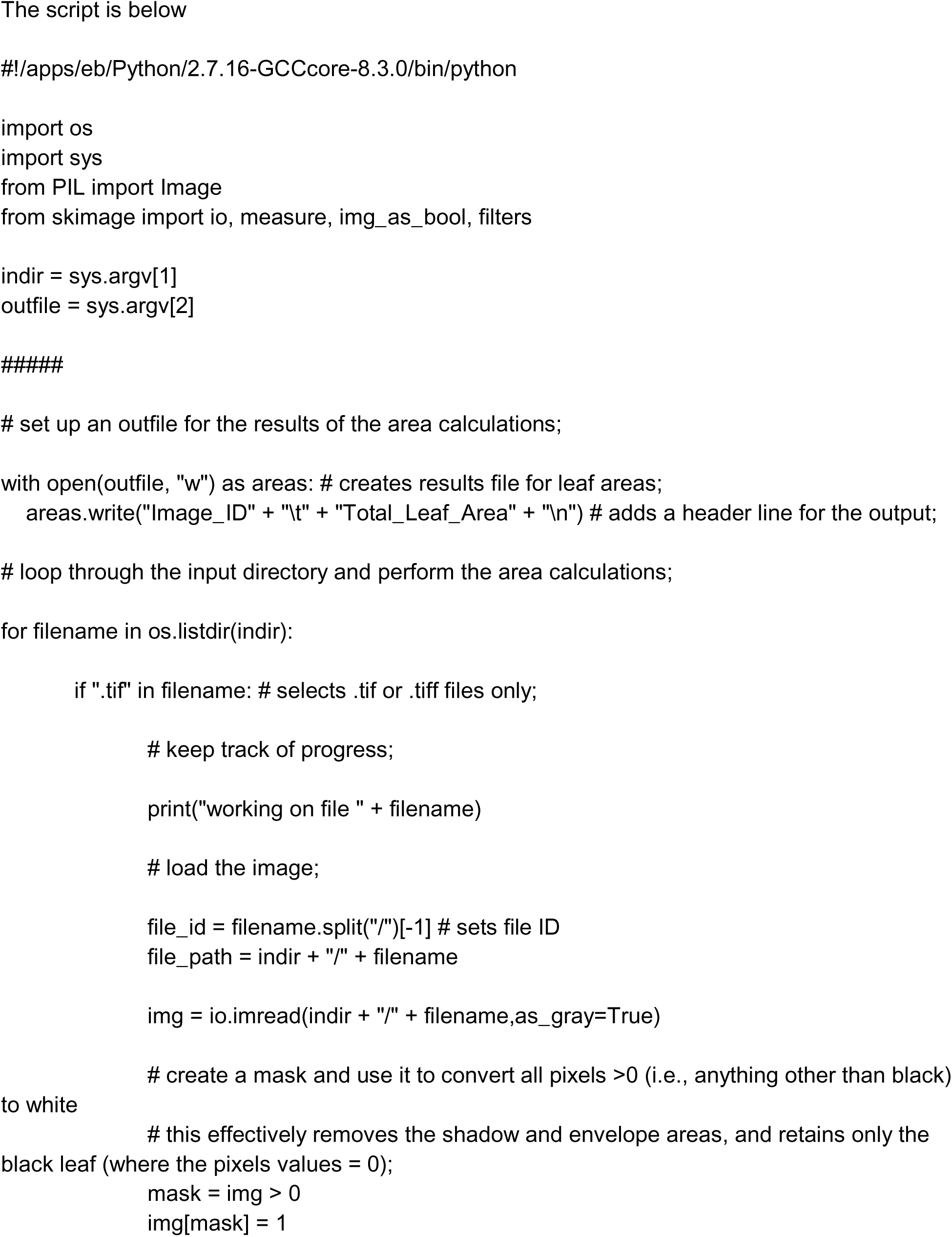
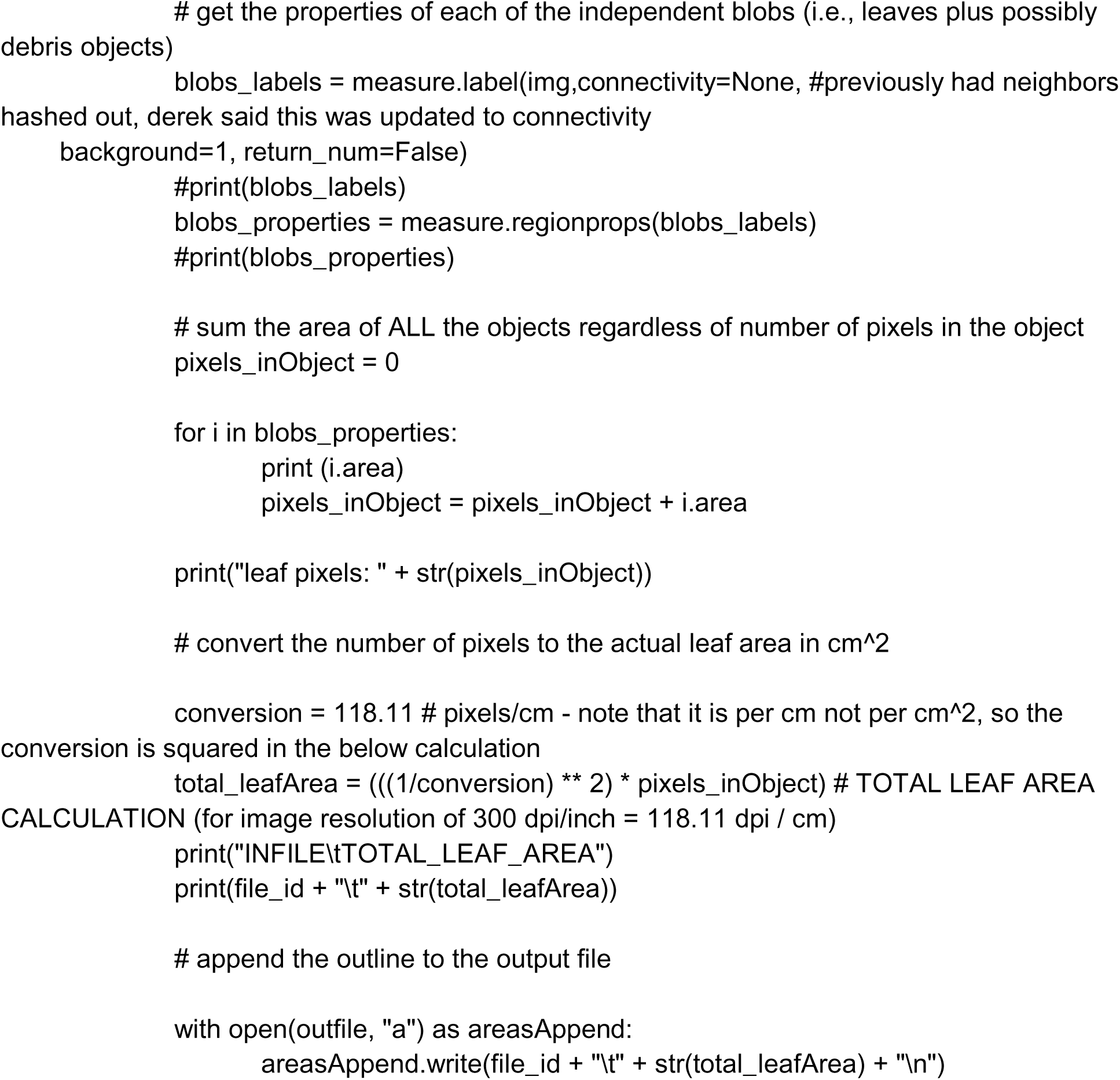

**Supplementary File 2**: Descriptions of phenotypes observed in the M0, M1, and M2 generations:

**Summary:** Irradiation treatment induced diverse phenotypes across the M0, M1 and M2 generations. There was a widespread recovery of initial responses, emergence of novel phenotypes in subsequent generations and limited cases of potentially heritable variants, where phenotypes persisted across generations.

**M0:** Initial irradiation responses were seen in the M0 generation, though lacking for the 5 Gy treatment.

- Each of the lines in the 10 Gy and 30 Gy treatment exhibited leaf phenotypes, including patches of light green tissue, wrinkled leaves and abnormally light or dark leaf coloration.
- We observed a range of malformed flowers, where the flowers were small, thin and narrow, or split down the side.
- Treatment-specific patterns emerged, where 5 Gys had only 5 lines with phenotypic responses. In addition, 30 Gy had an abundance of floral related phenotypes.

**M1:** The M1 generation was marked by recovery (absence of abnormal phenotypes) as well as novel phenotypes not observed in the M0 generation.

- The widespread leaf phenotypes were not observed as commonly as the M1 generation
- There was an increase in flowering related phenotypes as well as germination phenotypes, and meristem abnormalities.
- Interestingly, the new phenotypes were concentrated in the 5 Gy treatment (246 lines) and 10 Gy treatment (230 lines) compared to the 30 Gy treatment (30 lines)
- A small number of lines had flowering related phenotypes that were also observed in the M0 generation (5 Gy: 2 lines, 10 Gy: 3 lines, 30 Gy: 40 lines)

**M2:** The M2 generation had continued recovery and emergence of new phenotypes, along with lines across irradiation treatments exhibiting persistent phenotypes across generations.

- By the M2 generation, extensive phenotypic recovery occurred across all treatments, with a total of 446 lines no longer displaying abnormalities observed in M1 (5 Gy: 198; 10 Gy: 191 lines; 30 Gy: 57 lines).
- New phenotypes continued to emerge but at reduced frequency compared to M1, appearing in 74 lines (5 Gy: 32 lines; 10 Gy: 31 lines; 30 Gy: 11 lines).
- Persistent phenotypes, predominantly floral, were observed in 111 lines distributed relatively evenly across treatments (5 Gy: 45 lines; 10 Gy: 32 lines; 30 Gy: 34 lines).

